# Unleashing Precision and Freedom in Optical Manipulation: Software-Assisted Real-Time Precision Opto-Control of Intracellular Molecular Activities and Cell Functions

**DOI:** 10.1101/2024.02.09.579709

**Authors:** Bin Dong, R. Mike Everly, Shivam Mahapatra, Mark S. Carlsen, Seohee Ma, Chi Zhang

## Abstract

The traditional method in biological science to regulate cell functions often employs chemical interventions, which commonly lack precision in space and time. While optical manipulation offers superior spatial precision, existing technologies are constrained by limitations in flexibility, accuracy, and response time. Here, we present an adaptable and interactive optical manipulation platform that integrates laser scanning, chemical sensing, synchronized multi-laser control, adaptable target selection, flexible decision-making, and real-time monitoring of sample responses. This software-assisted real-time precision opto-control (S-RPOC) platform facilitates automatic target selection driven by optical signals while permitting user-defined manual delineation. It allows the treatment of mobile or stationary targets with varying laser dosages and wavelengths simultaneously at diffraction-limited spatial precision and optimal accuracy. Significantly, S-RPOC showcases versatile capabilities including adaptive photobleaching, comprehensive quantification of protein dynamics, selective organelle perturbation, control of cell division, and manipulation of individual cell behaviors within a population. With its unprecedented spatiotemporal precision and adaptable decision-making, S-RPOC holds the potential for extensive applications in biological science.

## Introduction

Optical imaging serves as a powerful tool for unraveling intricate biological processes. However, it cannot actively manipulate molecular targets and cellular behaviors. Active control of intracellular targets is crucial for deciphering correlations between biomolecules and cellular responses. Conventional approaches to manipulating molecular activities, such as chemical treatments or genetic interventions, lack spatiotemporal precision, potentially resulting in unintended side effects on non-targeted sites. Hence, to comprehensively understand dynamic molecular mechanisms and the causalities underlying cell behaviors, it is imperative to manipulate cellular targets with high spatiotemporal accuracy.

Laser-based optical manipulation methods leverage various light-matter interactions to modulate chemical processes at exceptional spatiotemporal precision. These techniques harness lasers to apply mechanical forces^1,2^, induce thermal perturbation^3^, regulate chemical processes^4^, activate proteins^5,6,^ generate reactive oxygen species^7,8^, and control photoswitchable compounds that interrupt molecular processes^9-11^. One paradigm of optical manipulation techniques relies on tightly focused laser beams that precisely trigger desired effects exclusively at the focal point. For example, optical tweezers use optical gradient forces to manipulate specific targets^1,2^. Focused pulsed lasers facilitate tasks such as cutting cell membranes for drug delivery^12,13^, disrupting mitochondria functions^14,15,^ and inducing cell fusion^16,17^. However, existing manipulation platforms rely on manually delineating targets based on prior knowledge from the sample, lacking the ability for chemical-based automatic target selection, and are unsuitable for highly mobile cellular entities. Alternatively, another approach of optical manipulation involves patterned illumination, utilizing widefield illumination and spatial light modulation techniques^18-20^. This method enables the simultaneous regulation of multiple neurons with optogenetics^21,22.^ However, it also necessitates prior sample information followed by the computation of a mask for spatial light modulation. The resulting light patterns often exhibit speckles, particularly in scenarios involving random or intricate patterns. Furthermore, it cannot simultaneously produce precise light patterns across diverse wavelengths. Compared with the focused laser approach, wide-field patterned illumination generally has diminished spatial accuracy and is incompatible with modalities that rely on tightly focused laser beams such as two-photon excitation fluorescence.

To address the limitations of existing optical manipulation technologies, we have developed a real-time precision opto-control (RPOC) technology that harnesses chemical signals to command separate laser beams for optical manipulation^11,23,24^. RPOC relies on a real-time optoelectronic feedback system, enabling automated selection of the laser interaction locus during scanning without the need for prior sample information. While the continuous-wave (CW) RPOC system is fully compatible with laser-scanning confocal fluorescence microscopes^24^, it does have limitations. First, the optical manipulation targets selected by the comparator circuit encompass the entire field of view (FOV), lacking the flexibility to delineate specific cells or subcellular areas for simultaneously creating different treatment conditions. Secondly, the determination of active pixels (APXs)^11,24^ solely relies on comparing sample fluorescence signals against a predetermined threshold within the comparator circuit. Consequently, targets that do not generate fluorescence signals cannot be selected by the system. Furthermore, the changes of fluorescence signals, caused by either photobleaching or augmentation by the action lasers, can influence APXs and the treatment laser dosage.

Here, we introduce a software-assisted real-time precision opto-control (S-RPOC) platform that delivers unprecedented flexibility, automation, and precision across chemical target selection, decision-making, laser control, optical manipulation, and real-time monitoring of cell responses. This platform orchestrates laser actions during scanning with exceptional selectivity, ensuring exclusive interactions with designated targets while leaving surrounding areas unaffected. S-RPOC integrates a dynamic decision-making subprogram enabling user-defined and optical-signal-determined selection of APXs, accommodating diverse configurations irrespective of prior knowledge. It also facilitates the concurrent creation of multiple manipulation conditions and enables comprehensive short- and long-term cell response monitoring during and post-optical manipulation. Applying S-RPOC, we demonstrate adaptable and precise photobleaching, achieve improved study of protein dynamics, generate reactive oxygen species (ROS) exclusively at designated targets, activate photoswitchable inhibitors in selected cells, and control cell division and viability via precise intracellular organelle manipulation. S-RPOC is compatible with all laser-scanning microscopes and elevates them to optical manipulation platforms with submicron precision and real-time responses. This advancement not only addresses the existing limitations in optical manipulation but also opens up new possibilities in biological science.

## Results

### Software-assisted real-time precision opto-control (S-RPOC)

The S-RPOC is integrated into a lab-built laser-scanning confocal microscope, as illustrated in **Figure 1a**. This platform is equipped with five continuous-wave (CW) solid-state lasers, ensuring coverage across excitation spectra of common fluorescent dyes and proteins. Specifically, the 405 and 532 nm lasers, commanded by acousto-optic modulators (AOMs), serve as the action lasers dictating cellular chemical processes. Other laser wavelengths are available for the excitation of fluorophores for chemical target selection or readout of cell responses. Three fluorescent detection channels are implemented in the confocal configuration to detect targets of interest and monitor cell responses. A stage-top incubator maintains the ideal culture conditions for cells and allows continuous monitoring of cell responses in the same field of view (FOV) for up to 72 hours.

**Figure 1.**
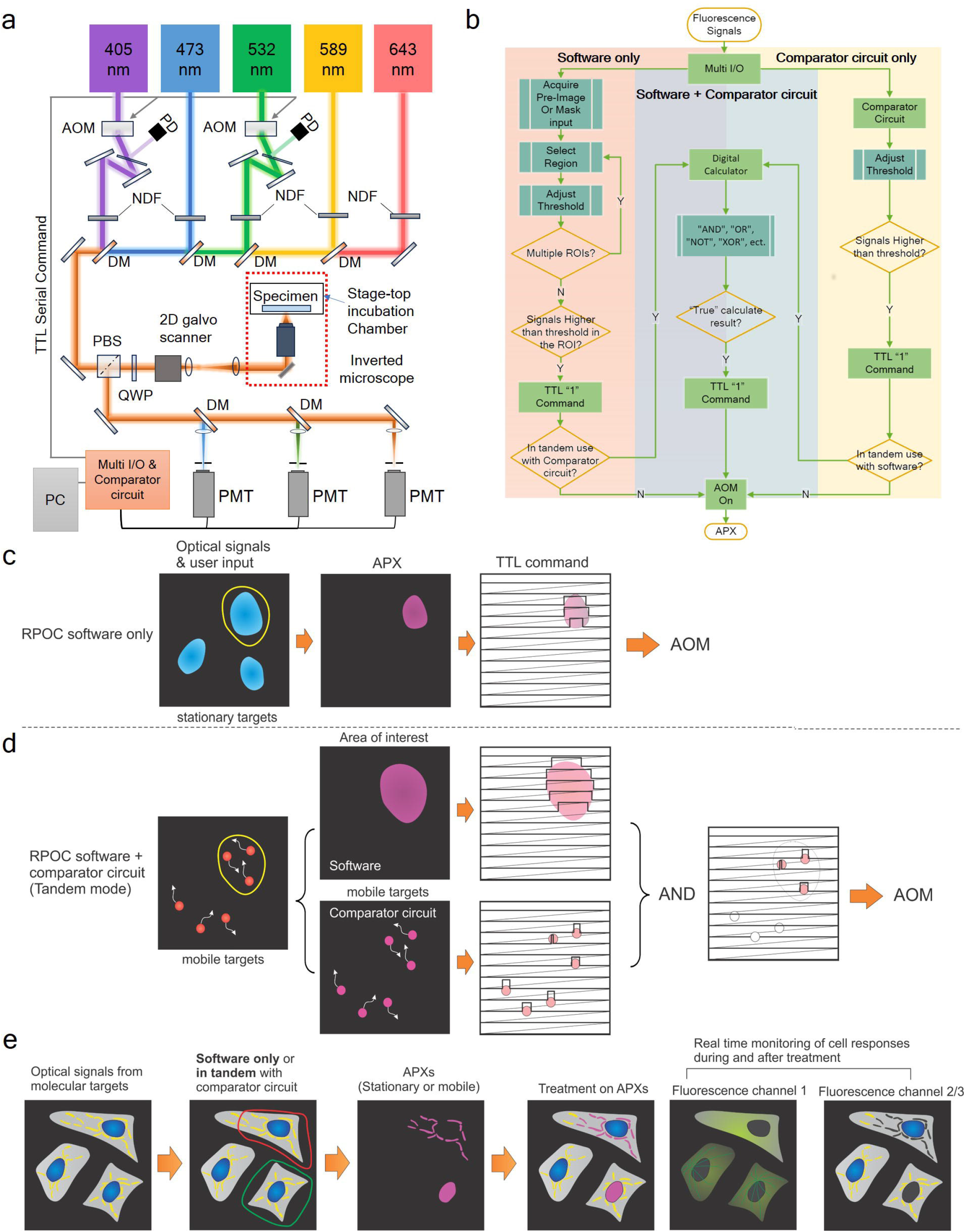
The S-RPOC technology overview. (a) The optical and electronic configurations of the S-RPOC setup. AOM, acousto-optic modulator; PD, photodiode; NDF, neutral density filter; DM, dichroic mirror; QWP, quarter-wave plate; PMT. Photomultiplier tube; PBS, polarizing beam splitter. (b) A schematic depicting the decision-making process across various S-RPOC operational modes. APXs, active pixels. (c) A schematic elucidating the generation of TTL control signal for AOM control, determined by optical signals and user input in the software-only mode. (d) The generation of TTL serial commands for AOM control in tandem mode involves applying an AND function to the area selection through the RPOC software and the mobile target tracking facilitated by the comparator circuit. (e) A schematic illustrating the action flow of S-RPOC. The yellow, blue, and green colors represent different fluorescence signals from the sample. The magenta color represents APXs. The red and green loops depict manual selection outlines for different action lasers utilizing the S-RPOC software.

The core opto-control processors of S-RPOC include a multifunction Input/Output (Multi-I/O) system and a multi-channel comparator circuit (**Figure S1**), operational either independently or in tandem. S-RPOC offers diverse methods for APX selection, allowing manual, automatic, or combined approaches. The decision-making flowchart is visualized in **Figure 1b**. Through the software subprogram, users can manually define areas of interest, followed by automatic target selection within the outlined areas. To perform this function, obtaining a sample image assists in delineating any region within the field of view (FOV) through an interactive software interface linked to the Multi-I/O (**Supplementary Video 1**). Within the selected area, intensity thresholding automatically identifies targets of interest (**Supplementary Video 2**). Different FOVs can be assigned to different action lasers for simultaneous optical manipulation. Furthermore, the software enables direct input of an image mask or a manually selected portion of the mask to choose APXs (**Supplementary Video 3**). These designated APXs can then be assigned to the action lasers, facilitating precise optical interactions with the targets. Notably, the manual area selection function facilitated by the software interface can be used in tandem with the comparator circuit (**Figure S2**). The circuit allows real-time comparison of the optical signals with a predetermined intensity threshold for APX selection. This integration offers an advantage for simultaneously tracking and precise control of moving targets within any user-defined area (**Supplementary Video 4**). The software-generated digital signals are computed with the TTL signals produced by the comparator circuit through built-in digital logic calculators including “AND”, “OR”, “NOT”, “XOR”, etc.

**Figure 1c** explains how the TTL command is determined from optical signals to control the AOM in the software-only mode. User input, such as adjusting the intensity threshold or uploading a mask, specifies the APXs that are synchronized with galvo scanning to send TTL ‘1’ to AOM only at APXs. In tandem mode, as illustrated in **Figure 1d**, the software-selected area of interest and the comparator circuit-selected APXs are computed with an AND function to exclusively address mobile targets within the chosen area. **Figure 1e** demonstrates the action flowchart employing S-RPOC for target selection and simultaneous monitoring of both APXs and cell responses. A comprehensive discussion of the functionalities of the platform, as well as the advantages and limitations associated with various modes, can be found in the **Supplementary Information** (**Note 1 and 2, Figure S3**). In summary, the software-only mode offers fixed laser dosage for each condition and mask input, albeit requiring prior knowledge of the sample and being applicable primarily for less mobile targets. Conversely, the comparator + software tandem mode enables the tracking and control of highly mobile targets (**Figure S4**), yet the laser dosage might be affected by alterations in optical signals. Both modes facilitate manual outlining of areas in flexible shapes and sizes, as well as automatic target selection based on optical signals. They also enable the selection of specific targets and the creation of diverse optical manipulation conditions within the same FOV. This capability represents a significant advantage of S-RPOC over traditional RPOC methodologies.

**Figures 2a-c** illustrate manual outlining areas on fluorescent microspheres for treatment with 405 nm and 532 nm lasers using a digital sketch board connected to the S-RPOC interface. The APXs on particles can be fine-tuned by adjusting intensity thresholds (**Supplementary Video 1, Figures S5,6**). The photobleaching effect, induced solely by the 405 nm laser, can be monitored both during and after laser treatment. **Figures 2d-f** show the photobleaching of fluorescent microspheres using a Purdue logo as the input mask. Additionally, **Figures 2g-i** display the bleaching of ‘The Mona Lisa’ and the inverted pattern on fluorescent microspheres using the painting mask and its contrast inversion (**Supplementary Video 2**). The execution of manual area selection or mask loading for S-RPOC takes only seconds. The delineated areas, each with separately adjustable APX thresholds, can be continuously added to different laser channels. These results highlight the adaptability of S-RPOC in controlling lasers for optical manipulation across any patterns and laser combinations.

**Figure 2.**
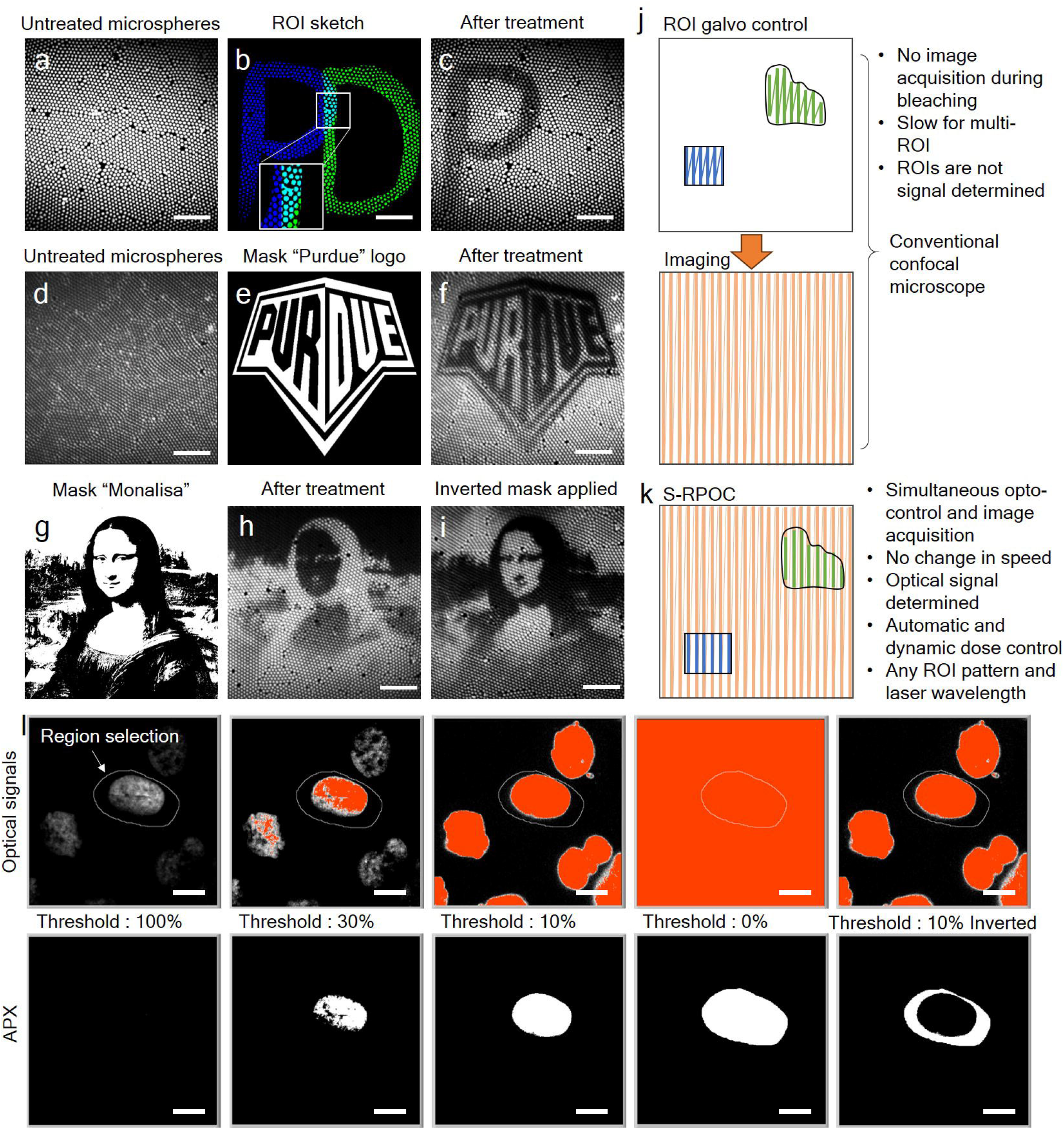
Flexible control of lasers and determination of APXs. (a) A fluorescence imaging of untreated 1 μm fluorescent polystyrene microspheres. (b) Manually designated areas for 405 nm (blue) and 532 nm (green) laser treatment, with the overlapped area shown in cyan and magnified for clarity. (c) The 405 nm laser treatment induces photobleaching in the selected area. (d-f) Photobleaching of a Purdue logo on fluorescent microspheres using the mask presented in panel (e). (g) The mask of the ‘Mona Lisa’ used for photobleaching. (h,i) Photobleaching of fluorescent microspheres using the mask in panel (g) and its contrast-inversed counterpart. (j) In a conventional confocal fluorescence microscope, the manually selected region of interest (ROI) for laser treatment is performed before imaging. (k) In RPOC, the APXs are automatically selected and guided by user input. Laser scanning, treatment, and imaging are performed simultaneously. (l) Examples of selecting APXs using different intensity thresholds in an outlined area encompassing a cell nucleus. The circle in the optical signal channel is manually delineated using the S-RPOC software interface. Scale bars: 10 μm.

**Figures 2j,k** describe the key distinctions between S-RPOC and conventional laser beam control. Different from Region of Interest (ROI) selection in a conventional confocal fluorescence microscope, where lasers scan through various ROIs before imaging, S-RPOC selectively activates or deactivates the action lasers at chosen APXs during imaging. The executions of image acquisition and opto-control occur simultaneous without interruption. Furthermore, S-RPOC maintains a consistent speed of optical manipulation and imaging regardless of the complexity of the treatment pattern. It also enables the simultaneous treatment of diverse areas in any pattern or distribution using different action lasers. Additionally, as shown in **Figure 2l**, in contrast to ROI selections in a standard confocal microscope, APXs in S-RPOC can be automatically chosen based on the intensity thresholding of optical signals from the sample. These APXs accurately align with desired molecular targets forming any distribution pattern and can be flexibly adjusted using the manual selection and intensity thresholding functions.

### Concurrently creating diverse optical manipulation conditions

One of S-RPOC’s major strengths lies in its capability to generate diverse optical manipulation conditions within the same FOV, enabling simultaneous comparisons of multiple parameters and enhancing manipulation throughput as exemplified in **Figures 3a-c**. The nuclei of HeLa Kyoto Histone-H2B-mCherry EGFP-Alpha-Tubulin cells were treated using a 405 nm laser at varying dosages, showcasing S-RPOC’s ability to compare different laser dosages and quantify changes within the same FOV in both mCherry-H2B and EGFP-Tubulin signals. The mCherry signal decrease observed in nuclei (**Figures 3d,e**) results from the generation of ROS induced by the 405 nm laser. A greater laser dosage leads to increased ROS generation, giving a faster fluorescence signal decay. Furthermore, S-RPOC permits simultaneous treatment of nuclei using 405 nm and 532 nm lasers within the same FOV (**Figures 3f-i**). In contrast to ROS, the 532 nm laser primarily induces photobleaching of mCherry-H2B signals, resulting in a distinct signal decay pattern (**Figure 3j**).

**Figure 3.**
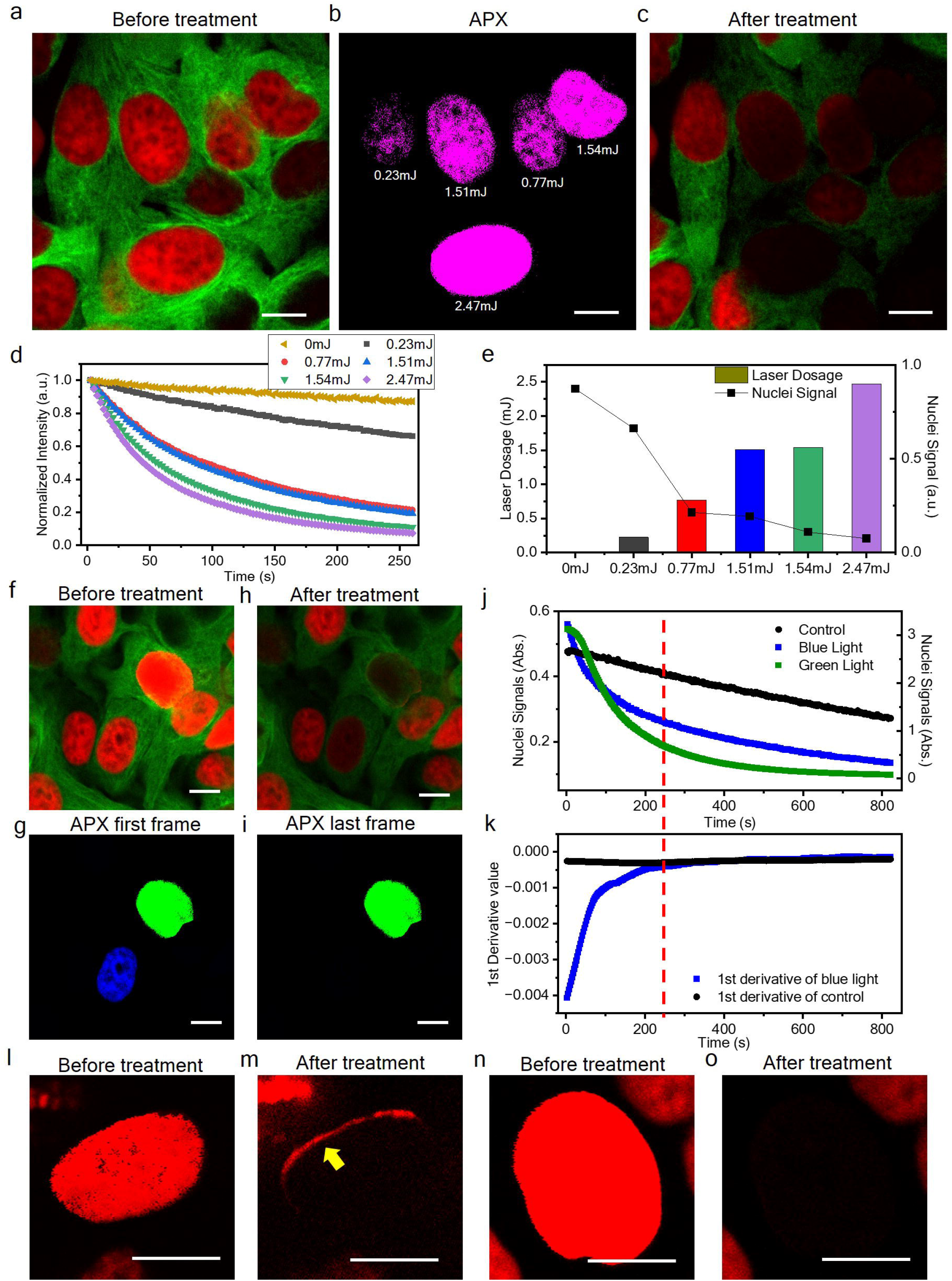
Simultaneous creation of various optical manipulation conditions in the same FOV. (a) A fluorescent image of HeLa EGFP-Alpha-Tubulin Histone-H2B-mCherry cells before treatment. The mCherry signals are displayed in red and EGFP signals are displayed in green. (b) APXs of 405 nm laser give different dosages to different nuclei. For each nucleus, manual delineation is performed followed by separate adjustment of the intensity threshold in each selected region. (c) The mCherry and EGFP signals after 260 s treatment. (d) The mCherry-H2B signal decay as a function of treatment time. (e) The correlation between laser dosage and the mCherry-H2B signals after treatment. (f-i) Similar to panels (a)-(c) but simultaneously treated one nucleus with a 405 nm laser and another with 532 nm lasers. The 405 nm laser control is performed in tandem with the comparator circuit box, resulting in an automatic stop of laser interaction after the signal decreases to the preset threshold. The 532 nm laser is solely operated by the software and thus maintains a consistent dosage during treatment. (j) The mCherry-H2B signal changes for the control, 405 nm (blue), and 532 nm (green) laser treatment of the cells in panel (f). (k) The first-order derivative of the signals in panel (j). The red dash line across j and k indicates the approximate stop point of the blue light treatment, which is around 250 s. (l,m) The mCherry-H2B signals before and after treatment using solely the software. The movement of the cell nucleus during treatment results in an untreated edge, as marked by the yellow arrow. (n,o) The mCherry-H2B signals before and after treatment using the software + comparator circuit tandem model. Object motion-induced photobleaching artifact is prevented. Scale bars: 10 μm

In **Figures 3f-i**, the combined use of the comparator circuit with manual nuclei selection is applied for the 405 nm treated nuclei. This approach enables the automatic cessation of optical treatment once the optical signal decreases to a predetermined level via dynamic determination of the APX through the comparator circuit. **Figures 3j,k** illustrate the automatic stop of ROS generation induced by the 405 nm laser when the signal decreases below the intensity threshold. In **Figures 3f-i**, the intensity threshold for the 405 nm treated nucleus is set at 50% of the initial fluorescence signal level. In the absence of the 405 nm action laser, a slow decline in fluorescence signal observed in the control group is caused by photobleaching induced by the 473 nm and 589 nm excitation lasers. The exertion of the 405 nm laser triggers a significantly accelerated signal decay owing to the generation of ROS. By comparing the first derivative of the signal decays, we can evaluate that a total laser dosage of 0.24 mJ over 250 seconds induces about 50% photodamage to mCherry molecules before the automatic stop of photo-interaction. Conversely, in **Figures 3f-i**, the treatment of nuclei with the 532 nm laser is solely managed by the software, employing a consistent laser dosage throughout the treatment duration without automated laser suspension.

Furthermore, the tandem use of the comparator circuit and software ensures precise treatment of moving targets within the designated area. Solely employing the software maintains constant APXs and laser dosages at the selected regions throughout the treatment, resulting in the walk-off of the object from APX due to target movement. Such a walk-off can lead to untreated areas of a moving nucleus, as illustrated in **Figures 3l,m**. This disparity is resolved when the comparator circuit is utilized in conjunction with the software, eliminating the untreated nucleus boundary (**Figures 3n,o**).

### Facilitating adaptable FRAP and FLIP for improved understanding of intracellular protein dynamics

Fluorescence recovery after photobleaching (FRAP) and fluorescence loss in photobleaching (FLIP) are established techniques to investigate the dynamics of fluorophores in live cells^25-27^. However, existing confocal fluorescence platforms used for FRAP or FLIP are constrained by limitations related to ROI selection and laser control. As illustrated in **Figures 2j,k**, conventional methods direct lasers to scan through manually chosen ROIs before imaging. This hinders the capturing of fluorophore dynamics information during the photobleaching process. Additionally, this traditional photobleaching approach is impractical in tracking complex patterns created by random fluorophore distributions. Furthermore, the separation of photobleaching and imaging processes prevents laser interaction with mobile fluorescent species. S-RPOC overcomes these constraints by employing optical-signal-driven real-time target selection and action lasers activation during laser scanning.

Using S-RPOC, we investigated protein dynamics in HeLa cells expressing Histone-H2B-mCherry and EGFP-Alpha-Tubulin. By outlining portions of two nuclei and adjusting intensity thresholds (**Figure 4a**), we selected APXs for 405 nm (240 μW) and 532 nm (320 μW) lasers. We tracked H2B-mCherry signals before, during, and after a 260-second optical treatment, observing distinct responses in treated and untreated areas within nuclei (**Figure 4b**). The 532 nm laser induced photobleaching of H2B-mCherry, while the 405 nm laser generated ROS, resulting in varied signal decrease patterns (**Figures 4a,b**) similar to **Figures 3g-k**. After treatment, the nucleus area treated with the 532 nm laser exhibited FRAP while the 405 nm treated area, due to the disturbance from ROS, continued to show a gradual signal decline similar to the control (**Figure 4c**). Further discussion is available in the **Supplementary Information** (**Note 5**). Monitoring mCherry-H2B signal changes in the treated area over 5 hours (**Figure 4d**), we observed continuous signal recovery in treated areas, in contrast to the initial decrease observed in untreated areas. This suggests that immediately after treatment, H2B loss due to protein mobility in the treated areas surpasses cellular H2B synthesis. Two hours post-treatment, cellular production of H2B overtakes migration-induced signal loss. This interplay between protein mobility and synthesis is also evident in the H2B signal increase rate from treated areas.

**Figure 4.**
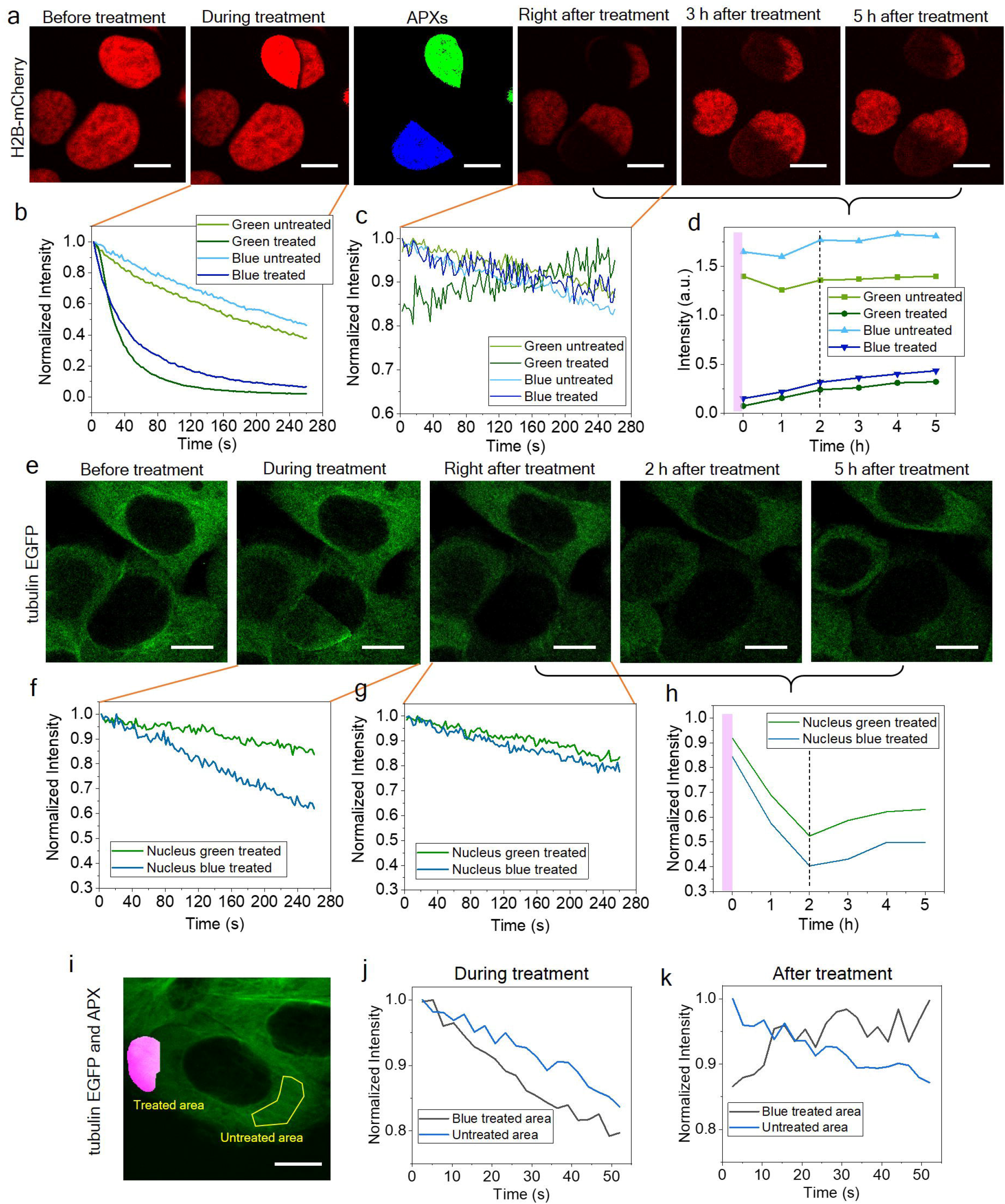
S-RPOC enables adaptive FRAP and FLIP. (a) mCherry-H2B signals (red) in live HeLa cells before, during, and after treatment, and APXs for 405 nm (blue) and 532 nm (green) lasers. (b) Changes in mCherry-H2B signals at treated and untreated nucleus sections during treatment. (c) Normalized mCherry-H2B signal changes at treated and untreated nucleus sections measured immediately post-treatment. (d) mCherry-H2B signal alterations at treated and untreated nucleus segments from 0 to 5 hours post-treatment. (e) The EGFP-tubulin signals in the same FOV of panel (a) before, during, and after treatment. (f-h) Similar to panels (b)-(d), exhibiting tubulin signals outside the nuclei of the treated cells. (i) EGFP-tubulin signals (green), APXs for 405 nm laser treatment (magenta), and the selected untreated region (yellow) for signal analysis. (j) EGFP-tubulin signal changes from the treated and untreated areas in panel (i) during treatment. (k) EGFP-tubulin signal change from the same areas immediately after treatment. Scale bars: 10 μm.

During the treatment of nuclei, we simultaneously monitored tubulin EGFP signals (**Figure 4e**). Our findings revealed that the 405 nm treatment of a partial nucleus led to a faster decrease in tubulin EGFP signals compared to the case of the 532 nm treatment (**Figure 4f**). This difference further supports the fact that the alterations induced by the 405 nm laser are attributed to ROS that oxidizes EGFP. After the treatment, the accelerated tubulin signal loss induced by ROS prominently slowed down (**Figure 4g**). Long-term monitoring displayed a two-hour continuous intensity decrease in tubulin-EGFP signals before recovery (**Figure 4h**).

The H2B protein exhibits a recovery half-time significantly longer than 5 hours (**Figure 4d**). To investigate proteins with a shorter diffusion time scale, we conducted a FRAP study of tubulin. APXs are selected at a subcellular location and the 405 nm laser was used for photobleaching (**Figure 4i**). The APX area demonstrates a faster EGFP signal decrease compared to the untreated area during RPOC (**Figure 4j**). After bleaching, the treated area shows a signal recovery within 12 seconds.

These results emphasize the power of S-RPOC in adaptive ROI selection and automated APX determination. This technology facilitates simultaneous monitoring of cellular responses during both treatment and recovery phases. Collectively, these capabilities enhance the FRAP and FLIP studies, offering improved insights into protein dynamics across various treatment conditions in a single FOV.

### Precisely perturbing mitochondria within specific cells in a population

Mitochondria act as cellular powerhouses. Light-induced ROS generated within mitochondria can disrupt mitochondrial functions, causing significant functional damage to cells^28,29^. S-RPOC enables the selective disruption of mitochondria inside selected cells within a population and the simultaneous monitoring of cell responses.

In **Figure 5a**, we label the mitochondria of HeLa Kyto EB3-EGFP cells using MitoTracker Red and outline a specific cell for exclusive 405 nm laser interaction with its mitochondria. Using the MitoTracker signals and adjusting the intensity threshold allows the automatic selection of mitochondria within the outlined cell (**Figure 5a**). The application of a 240 μW 405nm laser induces the MitoTracker signals to leak into the EB3-EGFP (**Figure 5a**). After treatment, the 405 nm laser interaction with mitochondria induces a significant loss in EGFP signals within the treated cell due to ROS generation. Such an EGFP signal decrease induced by light-mitochondria interaction is more pronounced than by the light-nuclei interaction as shown in **Figures 3 and 4**. This suggests distinct light-induced ROS disruptions to cells through different organelles. On the contrary, the MitoTracker exhibits much less signal decrease (**Figures 5a-c**). This indicates the higher stability of MitoTracker dye during the 405 nm laser interaction and ROS perturbation. However, the leakage of MitoTracker into the cytosol, evidenced by increased MitoTracker intensity outside mitochondria (**Figure 5b,d**), signifies mitochondrial damage and the loss of mitochondrial membrane potential caused by ROS.

**Figure 5.**
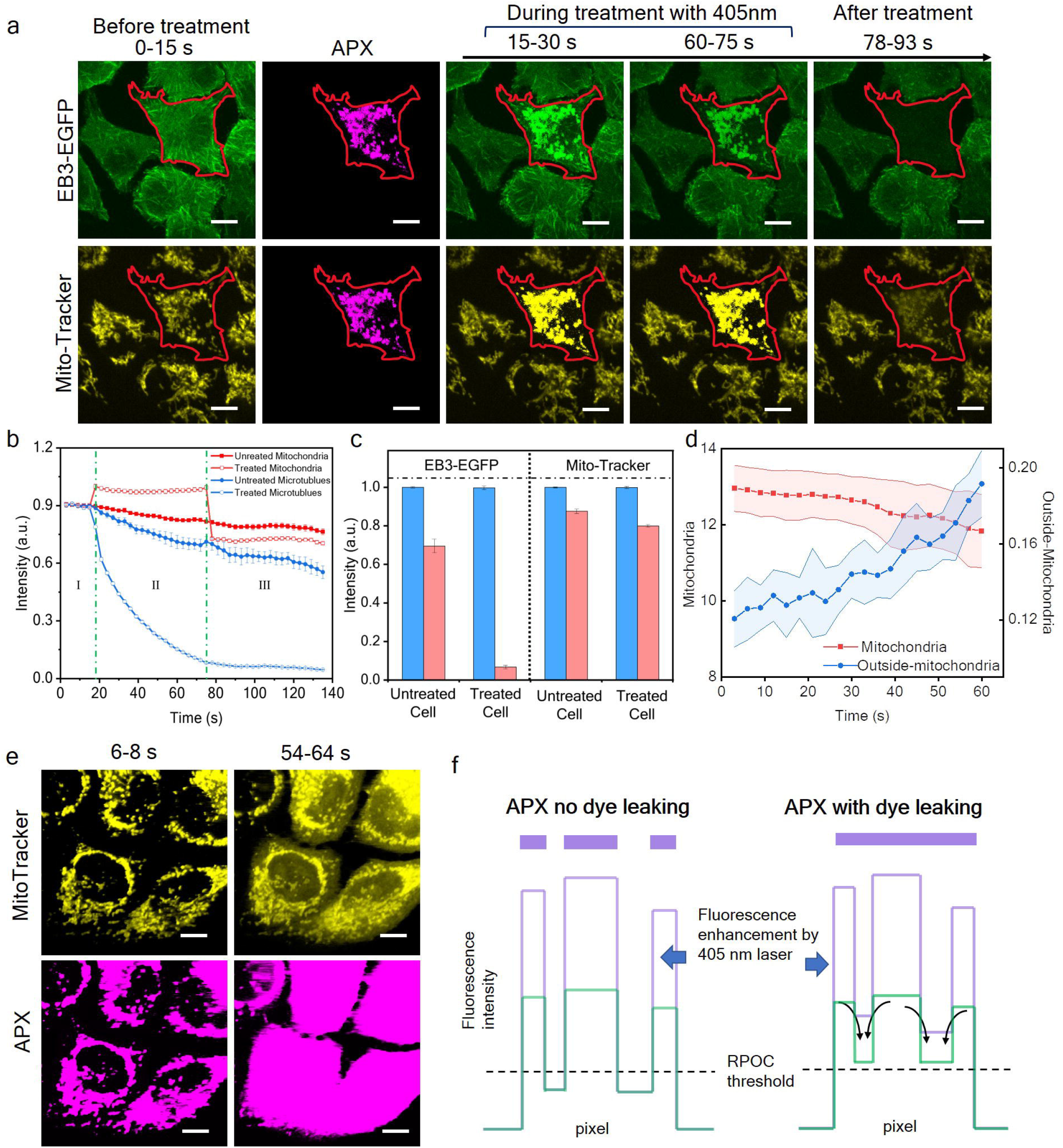
Inducing ROS selectively using a 405 nm laser at the mitochondria of a chosen cell. (a) EB3-EGFP (green) and MitoTracker (yellow) signals before, during, and after treatment of mitochondria by a 405 nm laser in a chosen cell. The APXs (magenta) are selected on mitochondria within the selected cell. The red outline represents the cell delineated using the S-RPOC software. (b) Fluorescence signals of EB3-EGFP and MitoTracker before (I), during (II), and after (III) treatment. (c) Intensity changes of microtubule and MitoTracker as a function of time before and after RPOC for both the treated and untreated cells. (d) MitoTracker signal changes post-treatment within and outside the mitochondria of the treated cell. (e) Time-lapse images of MitoTracker (yellow) and APX (magenta) during treatment when solely relying on the comparator circuit box showing the progressive disruption of APXs over time. (f) An illustration of APX disruption induced by the combined effect of signal enhancement by the 405 nm laser and MitoTracker leakage outside mitochondria. This disruption occurs during mitochondria treatment when only the comparator circuit is applied for real-time decision-making. Scale bars: 10 μm.

S-RPOC allows monitoring of critical changes in cellular energy flux and mitochondria function through EGFP and MitoTracker signals during and after blue light treatment of mitochondria using a consistent laser dosage. This accomplishment cannot be achieved with conventional RPOC solely with the comparator circuit. This limitation arises from the MitoTracker signal enhancement by the 405 nm laser and dye leakage into the cytosol, leading to a disruptive effect on APXs and laser dosage coordination as illustrated in **Figures 5e,f**.

### Regulating the behaviors of specific cells in both short-term and long-term scenarios

Apart from its ability to execute adaptive FRAP and FLIP and perturb mitochondria, S-RPOC holds the capacity to govern various short-term and long-term behaviors within a cell population through precise optical manipulation.

Together with photosensitive compounds, S-RPOC allows the inhibition of biochemical processes exclusively in selected cells within a population. For example, PST-1 is a photoswitchable microtubule inhibitor that can be activated by a blue laser and inactivated by a green laser^9^. Utilizing HeLa Kyoto EB3-EGFP cells for visualizing microtubule polymerizations, where the EB3 protein binds to the plus end of microtubules during polymerization^30,31,^ we employed S-RPOC to delineate a single cell for PST-1 activation (**Figures 6a,b**). The utilization of a 405 nm laser at 6 μW exclusively illuminated the outlined single cell resulting in the inhibition of microtubule polymerization only for this cell within the population, as shown in **Figures 6b,c**. Additional discussions can be found in **Supplementary Note 6**.

**Figure 6.**
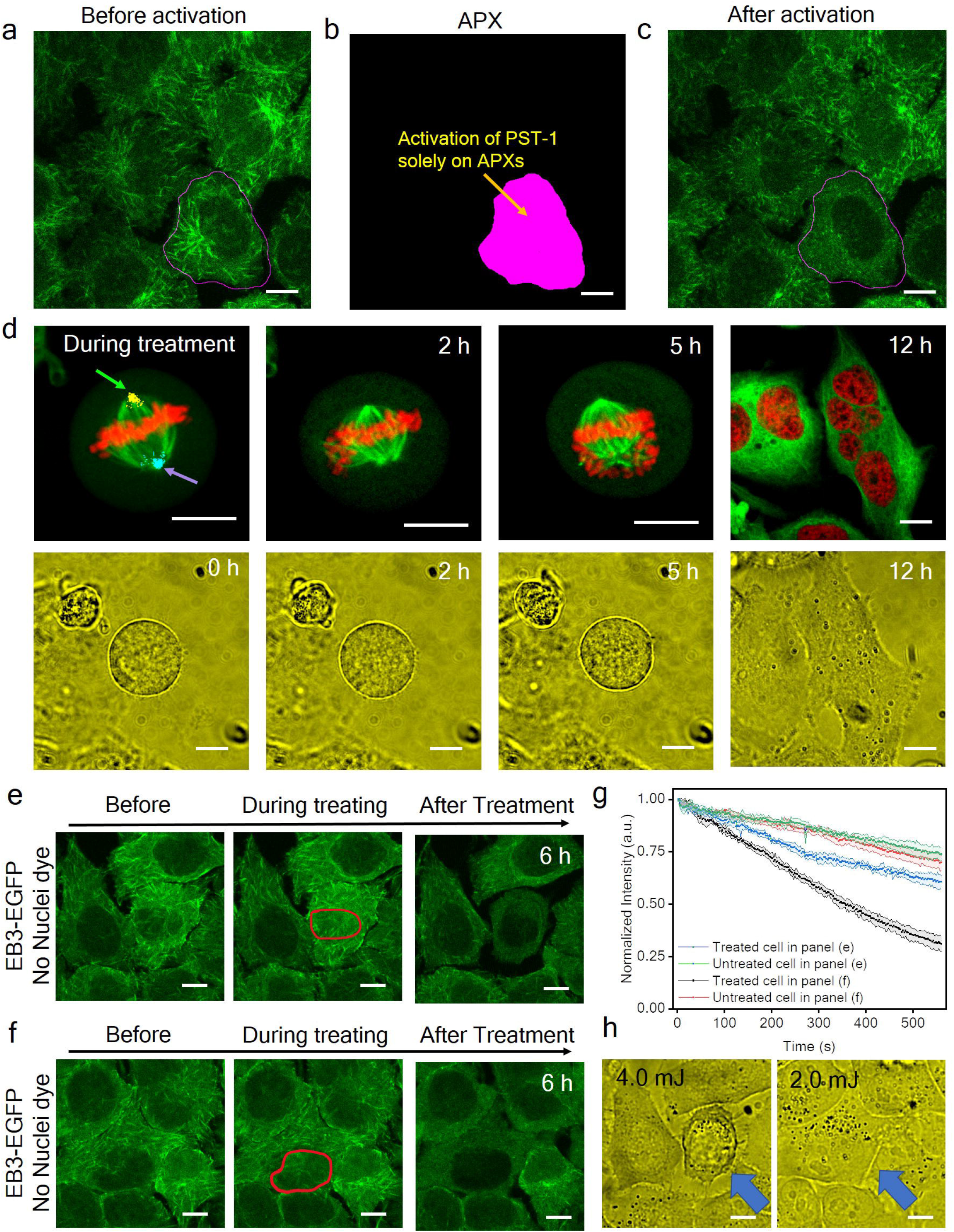
Regulating the short-term and long-term behaviors of specific cells. (a) EB3-EGFP signals from HeLa cells before PST-1 activation. The cell for treatment is outlined. (b) APXs selected for activating PST-1. (c) EB3-EGFP signals from HeLa cells after PST-1 activation. (d) Fluorescence signals of mCherry-H2B (red) and EGFP-tubulin (green) from HeLa cells at different time points and their corresponding bright field images. The two centrosomes are treated with 240 μW 405 nm laser (blue) and 240 μW 532 nm laser (yellow arrow). (e) Time-lapse images of HeLa cells expressing EB3-EGFP before, during, and 6 hours after treatment using the 405 nm for the circled area in the nucleus. The treatment time is 560 s and the laser dosage is 4 mJ. (f) Similar to panel (e) but the nucleus is treated for 280 s and a laser dosage of 2 mJ. (g) EB3-EGFP signal changes during treatment for cells in panels (e) and (f). (h) Bright-field images collected 10 hours after treatment from the same FOVs in panels (e) and (f) 10 hours after the treatment. Scale bars: 10 μm

S-RPOC demonstrates the capability to regulate cell division by employing blue-light perturbation specifically targeting the centrosome during the mitotic phase. The H2B-mCherry EGFP-tubulin HeLa cells are used to visualize both chromosomes and microtubules. The 532 nm and 405 nm lasers, both at 240 μW, are used to interact with two centrosomes of the cell (**Figure 5a**). Using S-RPOC, the APXs are selected only at the centrosomes by thresholding tubulin EGFP signals. The exposure with 0.38 mJ 405 nm laser at the centrosome disrupts the microtubule network in the anaphase, while the exposure with 0.41 mJ 532 nm does not change the spindle morphology (**Figures 5b,c**). Such a targeted optical perturbation of centrosomes by the 405 nm laser alters cell mitosis and results in the formation of multi-nuclei cells (**Figure 6d, Figures S11,12, Supplementary Note 7**). The entire process, starting from the treatment to the development of multinucleation, spans 15 hours and is continuously monitored within the same FOV aided by a stage-top incubator.

Utilizing the stage-top incubator facilitates the preservation of the cellular microenvironment, enabling the study of long-term cell responses. We conducted continuous monitoring of blue-light-induced cell perturbation, specifically targeting the nuclei, across different treatment durations and laser dosages. S-RPOC permits the selection of molecular targets or cellular compartments even without optical signals. Employing HeLa Kyoto EB3-EGFP cells, we outline nuclei areas in cells that show as voids in the EB3-EGFP signals. The results reveal that 405 nm laser treatment with a 2.0 mJ dosage directed at the nuclei causes a reversible perturbation to the cells, whereas a 4.0 mJ laser dosage results in irreversible apoptosis of the cells, as depicted in **Figures 6e-h and Figure S13**, and further discussed in **Supplementary Note 8**. Moreover, the targeted treatment of a cancer cell within a fibroblast coculture system using a 405 nm laser is also performed. (**Figure S14, Supplementary Note 9)**.

## Discussion

The S-RPOC technology developed herein is an advanced opto-control platform that enables real-time control of laser activities during scanning, thus granting unparalleled freedom in the optical manipulation of chemical processes within live cells. Its adaptive target selection, guided by optical signals and user input, allows the flexible selection of individual or multiple entities in cells, irrespective of their complexity and distribution. In addition, it offers simultaneous monitoring of cell responses during and after the manipulation, capturing both short-term and long-term changes using a broad array of readouts. S-RPOC stands out for its unparalleled versatility and capacity to manipulate solely the intended targets within live cells, leaving undesired locations unaffected. It can also concurrently establish multiple treatment conditions within the same FOV, thereby significantly reducing the overall experimental time. Furthermore, being able to compare the different treatment conditions in a single FOV minimizes variations induced by different cell populations. With S-RPOC, we demonstrated adaptive photobleaching of fluorophores, a better understanding of protein dynamics, inhibition of tubulin polymerization in selected cells, ROS generation only at targeted organelles within the cell of interest, and the control of cell division by centrosome stimulation. These diverse applications exemplify the extensive potential and broad impact of S-RPOC technology in biological sciences.

The spatial resolution and opto-control precision of S-RPOC are currently diffraction-limited, operating at a scale of about 300 nm. The laser scanning system has a fast-axis scan rate of 1 kHz, typically allowing a 0.2 to 1 Hz frame rate used in this study. The choice of optical manipulation lasers extends beyond the current 405 and 532 nm CW lasers. It encompasses a broad range of lasers such as ultraviolet, near-infrared, mid-infrared, and ultrafast lasers, allowing for a diverse range of opto-control applications. Beyond generating ROS and altering chemical states, these lasers can selectively induce heating^32^, generate localized plasma^33^, or perturb samples through various mechanisms, enabling the study of cellular responses to a broad spectrum of stimuli directed at specific organelles.

Beyond live mammalian cells, S-RPOC has the potential to extend its applications to precise manipulation of chemical targets in multicellular systems and larger model organisms such as cultured organoids, brain slices, and *Caenorhabditis elegans*. In microbiology, S-RPOC can be applied to track and manipulate highly mobile bacteria cells of interest without affecting unwanted locations. The site-specific control of chemical targets, facilitated by S-RPOC, also opens new possibilities in directing stem cell differentiation and modulating embryo development.

## Methods

### The S-RPOC platform

The schematic of the S-RPOC platform is illustrated in **Figure 1a**. This system incorporates five continuous-wave (CW) solid-state lasers (405 nm, 473 nm, 532 nm, 589 nm, 643 nm, CNI Laser). The output power of the lasers can be adjusted via variable neutral density filters (NDF, 54-081, Edmund Optics) in their respective beam paths. The 405nm and 532nm lasers, designated for optical manipulation, are controlled separately by two acousto-optic modulators (AOMs, M1205-T80L-1 with 552F-2 driver, Isomet). After the AOMs, the first-order diffractions of these lasers are combined collinearly with other lasers using long-pass dichroic beam splitters (DM, #69-887, #69-888, #69-889, #69-890, Edmund Optics). A ‘1’ TTL command deflects the action lasers to the target while a ‘0’ TTL command blocks the action laser. Microscope coverslips which are placed after the AOMs reflect a small portion of the first-order beams into photodiodes (PDA10A2, Thorlabs) for direct visualization of the APXs of the 405 nm and 532 nm lasers. The combined laser beams then pass through a polarizing beamsplitter (PBS251, Thorlabs) and a quarter-wave plate (10RP454-1B, Newport) before entering a 2D galvo scanner (Saturn-5, ScannerMax). This galvo scanner is integrated into an inverted microscope frame (IX73, Olympus) along with a 3D translational stage (H117 with Motor Focus Drive and ProScan III system, Prior) to accommodate the sample. A stage-top incubator (OTH-STXF-WSKMXCO2O2, Tokai Hit) is employed to maintain the CO_2_ level, temperature, and humidity, allowing prolonged opto-control and monitoring of cell responses within the same FOV. A camera (Omax 18MP Camera) is set up in transmission mode to capture bright-field images of cells. A water-dipping objective lens (UPlanSApo-S, 60X, NA = 1.20, Olympus) is utilized to focus laser beams onto the sample for RPOC and fluorescence imaging purposes.

The fluorescence signal detection operates in an epi-confocal mode. Fluorescence signals are directed towards three photomultiplier tubes (PMTs, H7422-40, Hamamatsu) via a polarizing beamsplitter and are separated by two long-pass dichroic beam splitters sequentially (FF552-Di02, FF648-Di01, Semrock). Pinholes (P300HK, Thorlabs) are positioned at the sample conjugate plane, aligned with the focal position of the lenses before the PMTs. Three bandpass filters are used for the three PMTs for signal detection (FF01-509/22, Semrock, ET642/80m, Chroma Technology Corporation, and FF01-680/42, Semrock). The PMT output currents across all channels are converted to voltage and amplified using three preamplifiers (PMT4V3, Advanced Research Instruments Corporation). The amplified signals are then routed to the opto-control units, which play a pivotal role in determining APXs for optical manipulation. The control units consist of a Multi-I/O system (PCIe-6363 paired with BNC-2120, National Instruments), a comparator circuit, and interactive LabVIEW software. Manual delineation of desired areas can be accomplished through an electronic sketch pad or mouse using the interactive software interface (**Supplementary Video 1**). The threshold for the TTL command can be adjusted both via the software and comparator circuit. The ultimate decision to activate the APXs can be made solely through the software, using the comparator circuit, or via logical calculations derived from both the software and the circuit. The lasers, optical detectors, and laser scanners are housed within a light-tight enclosure, allowing the system to operate in ambient room light.

To enable APX selection using the software, the synchronization between the Digital Output (DO) transmitting TTL commands and the Analog Output (AO) governing the galvo scanner is crucial. In the software-only mode, an acquired fluorescence image from the sample or a provided mask input is transformed into a TTL time series. This series is directed in real-time to the AOM of either the 405 nm, the 532 nm laser, or both, during laser scanning. A ‘1’ TTL command signifies an APX, precisely activating the laser at the targeted pixel, while a ‘0’ TTL command blocks the action laser. In the software + comparator circuit tandem mode, the TTL command selecting the area by the software is directed to one input of the comparator circuit. Simultaneously, the fluorescence signal from mobile targets is directed to another input of the comparator circuit to compare with the intensity threshold. This result is logically computed with the area selection TTL in real-time via AND, OR, or NOT functions (**Figure S2**). The final outputs after logic computation control the AOMs. The digital AND function determines APXs on mobile targets within the selected area, while the digital NOT allows users to assign APXs on mobile targets outside the selected area.

### The laser dosage calculation

When the comparator circuit is applied, the laser dosage is impacted by the changes in optical signals from the sample. To quantify the laser dosage received by a single cell, two photodiodes are applied to detect a small portion of 405 nm or 532 nm lasers deflected from the first-order output of the AOMs. Such detections generate APX images. To calculate laser dosage received by a single cell, manual delineation of an area encircling all the APXs in the selected cell is performed in ImageJ. The laser dosage is computed as

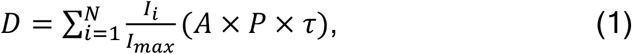

where A represents the size of a delineated area encompassing all APXs for a single cell, I_i_ is the average intensity within the delineated area for frame i, P denotes the laser power on the sample, τ is the pixel dwell time, and N stands for the total number of frames for laser treatment. The unit of the laser dosage is joule (J). I_*max*_ can be achieved by reducing the intensity threshold to zero using the comparator circuit. The image processing, region outlining, and quantifications are conducted using the built-in functions of Image-J (Fiji).

Alternatively, when only applying the software, both the APXs and laser dosage remain consistent throughout the treatment. Hence, the APX information can be obtained directly from the Multi-I/O. Except the above equation, another method to calculate the laser dose is using the equation

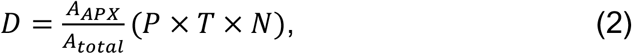

where A_*APX*_ and A_*total*_ are the total areas of the APXs and entire image, respectively, calculated using the built-in functions of Image-J (Fiji). P denotes the laser power on the sample, T is the total time of for each frame, and N is the total number of frames. Under the software-only condition, the laser dosage quantified using these two methods agrees with each other.

### Cell preparation

HeLa Kyoto EB3-EGFP cells and HeLa Kyoto EGFP-alpha-tubulin/H2B-mCherry cells are purchased from Biohippo. Panc.10.05 and CAF19 cells are obtained from Dr. Bumsoo Han’s research group at Purdue University. Cells are cultured in Dulbecco’s Modified Eagle Medium (DMEM, ATCC) with 10% fetal bovine serum (FBS, ATCC) and 1% penicillin/streptomycin (Thermofisher Scientific). The cells are seeded in 35 mm glass-bottom dishes (MatTek Life Sciences) with 2 mL culture medium and then incubated in a CO_2_ incubator set at 37 °C and 5% CO_2_ concentration. Upon reaching approximately 50-70% confluency, the cells are used for treatment or live-cell optical manipulation in the stage-top incubator.

### Fluorescent microspheres and MitoTracker labeling of cells

The 1 μm fluorescence polystyrene microspheres (Lot No. 2051) are purchased from Phosphorex Inc. Their excitation and emission maximums are 460 nm and 500 nm, respectively. The microsphere solutions are deposited onto a coverslip and air-dried to form an evenly distributed single-layer microsphere film for photobleaching experiments.

HeLa Kyoto EB3-EGFP cells are first seeded in 35 mm glass-bottom dishes and cultured overnight to reach a confluency of around 50-70%. Mitotracker Red CMXRos is added to the culture medium at a final concentration of 200 nM. The cells are then incubated for 30 min at 37 °C and 5% CO_2_ concentration before S-RPOC.

### PST-1 preparation and treatment

PST-1, synthesized as described in prior publications^11^, is dissolved in dimethylsulfoxide at a concentration of 2 mM to create the stock solution. Before cell treatment, the PST-1 stock solution undergoes exposure to a 532 nm laser (CNI laser) for 5 seconds to convert PST-1 into its trans-inactivated form. Following this, cells are treated with the inactivated PST-1 at a final concentration of 4 μM for 15 minutes before performing RPOC. The 405 nm laser in the S-RPOC system is utilized to selectively activate PST-1 exclusively at specified cells. The 405 nm laser power for PST-1 activation used in S-RPOC is 6 μW.

## Data availability

The data supporting the findings of this study are available within the paper and its supplementary information files. Additionally, supporting data may be obtained from the corresponding author upon reasonable request.

## Code availability

The source code developed for this research is currently undergoing a patent application process. We plan to make the code publicly available upon the conclusion of the patent application. Patent rights may apply.

## Acknowledgments

This work is supported by NIH R35GM147092. The authors acknowledge Dr. Mingji Dai and Dr. Yiyang Luo for the synthesis of PST-1, which was published previously. Additionally, we acknowledge Dr. Bumsoo Han for sharing the Panc.10.05 and CAF19 cells used in this study.

## Author contributions

C.Z and B.D. designed the experiment. B.D. and S.H.M. performed the experiments. R.M.E. designed and programmed the S-RPOC software. B.D. and R.M.E. debugged the S-RPOC software. S.H.M. and S.M. prepared cells, fluorescence labeling, and PST-1 treatment. M.C. designed and fabricated the comparator circuit. B.D. and C.Z. analyzed the results. C.Z. obtained funding for this research. C.Z. and B.D. wrote the manuscript.

## Supplementary Information

Supplementary Notes, Figures, and references are available.

