## Supplementary Figures and Notes for "Unleashing Precision and Freedom in Optical Manipulation: Software-Assisted Real-Time Precision Opto-Control of Intracellular Molecular Activities and Cell Functions"

### Supplementary Notes

#### Note 1. The comparator circuit box and its functions

The two-channel comparator circuit box used in this research is assembled with commonly available electronic components. It can operate independently or in tandem with the S-RPOC software, enabling real-time tracking and exclusive optical manipulation of mobile targets within the sample.

The ports and function switches of the comparator circuit box are shown in **Figure S1a**. It integrates two identical comparator circuits that can function separately or in combination. Each circuit includes an 'analog input' for receiving signals from an optical detector. The signals are compared with a manually adjustable preselected threshold via an adjustment knob or a digital threshold input. The resultant TTL signal after comparison can output directly from the 'direct comparator output' ports or perform logic computation with the TTL signal generated from the other circuit. Switching between the two modes is facilitated by signal path switches. In the digital logic mode, the 'direct comparator output' can also serve as a signal input. In addition, a buffered analog signal output is provided for each comparator circuit to display the optical image. Digital logic functions such as AND, OR, and NOT are available for computations between the two channels. Digital NOT functions are positioned before the 'direct comparator output' and 'digital logic output', enabling inversions of APXs if needed. Two 'digital logic output' copies are available post-digital logic computation, allowing simultaneously commanding AOMs and displaying APXs. Before each digital logic output, an 'on' and 'off' switch is available to facilitate sending '1' and '0' TTL signals to constantly turn on and off the connected AOMs.

When optical signals from two separate detection channels are linked to two analog inputs, the comparator circuit allows for selecting APXs based on criteria established separately by both comparators. Alternatively, when the optical signals from a single detection channel are divided and connected to both analog inputs, it permits the selection of APXs from the cellular entities exhibiting optical signals within an intensity passband. Either comparator circuit can also function independently without interference from the other. Regardless of the conditions, outputs from the two comparator circuits can independently control two AOMs, commanding different action lasers.

In the scenario where the comparator circuit is used in tandem with the S-RPOC software, its connection is illustrated in **Figure S2a.** One comparator circuit operates as previously described. The software TTL signal output can feed as the input from the 'direct output channel' of the other unconnected comparator circuit. The 'signal path switch' from the disconnected comparator circuit must be directed to the digital logic function. By selecting the AND function of any 'digital logic output', the comparator circuit will target mobile chemical entities in the area delineated by the S-RPOC software. Note that this 'in tandem' mode enables the treatment of a specific subset of targets with the treatment of another subset selected solely using the software (as shown in **Figure 1d**). To select mobile targets outside the delineated region, the connection is shown in **Figure S2b**. Note that if the 'invert' function in the bottom comparator circuit is disabled (set to non-invert), the connection in **Figure S2b** offers the same function as **Figure S2a**.

**Note 2. S-RPOC function modes**

S-RPOC can operate in two configurations: The software-only mode and the software + comparator circuit in tandem mode. The primary functional characteristics, as well as the advantages and disadvantages, are elucidated in **Figure S3**.

In the former configuration, the comparator circuit box is bypassed, and the digital output from the Multi I/O system is directly used to command AOMs. This modality is ideal for controlling less mobile molecular targets or when the optical signals from the sample are significantly affected by the treatment. Typically, an optical image is acquired to aid in selecting regions of interest (ROI). Laser dosage within regions, selected by the software based on the acquired image, remains consistent throughout the treatment process. Furthermore, it permits the input of a digital mask to guide laser interactions, as shown in **Figures 2d-i**. The software provides four digital outputs for users, allowing manual selection, digital mask input, or partial selection of the input mask separately for up to four action laser sources. Each channel includes independent intensity threshold functions aiding in the automatic selection of chemical targets based on optical signals. The mask input enables laser manipulation based on complex post-image analysis beyond intensity thresholding. For manual delineation of targets using a digital sketch pad or mouse, the selected area can be continuously added to each channel with distinct intensity thresholds for individual actions. These unique functionalities enable flexible user input, automated target selection based on chemical signals, laser dose control through intensity thresholding, and simultaneous control of multiple laser wavelengths. Various treatment conditions, including differences in area, laser dosage, and wavelength, can be simultaneously generated within a single Field of View (FOV). This capability facilitates effective comparison of diverse treatment conditions, significantly enhancing RPOC throughput. Furthermore, it permits the study of interactions among different cells treated in distinct manners.

The tandem configuration is particularly suitable for manipulating highly mobile targets in cells. In contrast to conventional RPOC, the S-RPOC tandem mode allows for precise control of moving targets within a sub-FOV with different lasers and comparisons of different treatment conditions in the same FOV. It additionally prevents imprecise laser treatments caused by slow target drifting, exemplified by the nucleus movement in **Figures 3l-o**. However, in this mode, the laser dosage can be affected by the changes in optical signal during treatment. One notable disadvantage is the difficulty in ensuring a constant laser dose over time. However, this adaptive and automatic APX adjustment permits dynamic photo interactions. The photobleaching or light-induced ROS generation can automatically stop when the optical signals fall below the intensity threshold established by the comparator circuit. For example, as demonstrated in **Figures 3f-k**, the interaction of the 405 nm laser with the nucleus automatically stops when the nucleus fluorescence signals drop to 50% of the initial signals.

The tandem configuration is particularly advantageous for instantaneously tracking and modulating laser interactions for highly mobile targets. In **Figure S4 and Video S4**, the EB3-EGFP signals from HeLa cells and the identified APXs utilizing the EB3 comets are presented for 532 nm and 405 nm laser. Through the S-RPOC software, individual cells can be delineated, allowing lasers to exclusively engage with EB3 comets in the respective cells in real-time. In **Figure S4**, the low power of the action lasers in both treatment conditions prevents noticeable photobleaching or disruption of EB3 comets through targeted actions. This figure highlights the ability to selectively choose APXs on highly dynamic molecular targets using different lasers in tandem mode.

### Note 3. S-RPOC target selection using the software and intensity thresholding

S-RPOC permits adaptable APX selection through both manual delineation and automated target selection based on optical signals. **Figure S5** gives examples of delineating a single ROI involving the molecular targets. Encircling a cell nucleus expressing mCherry-H2B signals and adjusting the intensity threshold within the software allows for the selection of APX covering different areas of the nucleus. Lowering the threshold level enables the selection of a larger area within the nucleus. Furthermore, reducing the threshold levels to the minimum allows for the selection of APX covering the entire selected area. In addition, using the invert function, APXs can be selected on pixels having optical intensity below the threshold within the delineated area. The ‘In Range’ function permits the selection of APX for entities falling within any intensity range.

**Figures S6a,b** demonstrate the selection of multiple sub-FOV areas and the simultaneous adjusting of the intensity threshold of all these areas using the ‘Multi Selection’ function. **Figures S6c-e** illustrate the sequential addition of APXs to the same treatment channel by outlining different areas within the image. The intensity threshold for the sub-FOVs in each action is individually adjustable. **Figure S6f** shows selecting the APXs using the invert function after sequentially adding all the sub-FOVs. **Figures S6g,h** show choosing APXs using manual delineation and intensity thresholding for treatment using different laser lines (Line 4: 405 nm laser; Line 5: 532 nm laser). **Figure S6i** displays the treated areas by blue and green laser lines within the same FOV. **Figure S6j** shows switching the green laser treatment APXs using the ‘invert’ function.

These examples exemplify the flexibility in selecting APXs for optical manipulation and adaptive control of laser doses at any target. **Supplementary Video 1** illustrates these functions aside from using intensity thresholds to adjust the total number of APXs, the laser dosage can also be controlled by adjusting the output power of the action lasers.

### Note 4. Laser dosage calculation for software-only and in tandem with comparator circuit.

The laser dose calculations differ when employing only the software as compared to incorporating the comparator circuit. Detailed equations and explanations can be found in the Methods section. When using only the software, the TTL command remains '1' throughout the entire APX, while outside APXs, the TTL command is consistently '0'. Consequently, the APX intensity has only two values. **Figure S7** illustrates the control of the green laser using this software-only mode for nucleus treatment.

When the comparator circuit is applied, either independently or in tandem mode, the optical signal is continuously compared with the threshold in real time. The fluctuations in optical signals may surpass the threshold for varying durations, resulting in diverse APX signals. In **Figure S7**, the blue laser operates in tandem mode for nucleus treatment.

#### Note 5. mCherry-H2B signal changes during and after treatment

In **Figure 3a-h**, S-RPOC facilitates comprehensive monitoring of the entire photo-interaction process with 405 nm and 532 nm lasers, followed by FLIP and FRAP assessments. During treatment, distinct signal decay patterns are observed in response to the blue and green lasers, induced by ROS and photobleaching, respectively. Immediately after treatment, the region treated with the green laser starts signal recovery, attributed to FRAP, while the area treated with the blue laser demonstrates continuous signal decay similar to the untreated case. The normalized signal decay curves shown in **Figure 3c** exhibit similar decay rates for all untreated areas and the blue laser-treated area. However, in absolute values, the signal decay in the untreated areas is more pronounced compared to both the decay in the blue-treated area and the rise in the green-treated area (**Figure S8**). The signal rise induced by the FRAP of mCherry in the treated region surpasses the impact of photobleaching, whereas the combined ROS+FRAP process does not surpass the photobleaching effect, leading to a slow signal decline. Strategies to mitigate the photobleaching effect include reducing the excitation laser power, decreasing the pixel dwell time, and enlarging the FOV.

#### Note 6. Fluorescence signal changes of EB3-EGFP HeLa cells

End-binding protein 3 (EB3) attaches to the plus end of microtubules. Therefore, EB3-EGFP signals, typically appearing as comets, allow us to visualize microtubule polymerization in live cells. The reduction in EB3-EGFP signals serves as a metric for quantifying the generation of ROS in cells [S1]. In **Figure S9**, EB3-EGFP signals excited with 25 μW 473 nm laser and a pixel dwell time of 10 microseconds show a gradual signal decrease due to the photobleaching of EGFP molecules by the 473 nm laser.

When MitoTracker Red is utilized as a label, mitochondria can be simultaneously visualized with EB3-EGFP signals from separate PMT channels. As shown in **Figure S10a**, the fluorescence signals from both molecular entities are excited by 25 μW 473 nm and 10 μW 589 nm lasers. Negligible EB3-EGFP signal decay is detected after 90 seconds of laser illumination.

When RPOC is performed targeting mitochondria using the conventional approach by employing only the comparator circuit box, the leakage of MitoTracker fluorescence into the EGFP channel due to 405 nm laser interaction and the diffusion of MitoTracker outside mitochondria are detected after 20 seconds of treatment using the 25 μW 405 nm laser (**Figure S10b**). The MitoTracker leakage and signal enhancement disrupts APXs, as explained in **Figure 5e**. In comparison, when S-RPOC is applied to maintain a consistent laser dose specifically at the mitochondria, as demonstrated in **Figure 5a**, such APX disruptions do not occur.

#### Note 7. Controlling cell division by the treatment of the centrosomes using a 405 nm laser

For regulating cell division by targeting centrosomes, S-RPOC is employed to select a confined area encompassing the centrosomes, indicated by the conjunctions of microtubule spindles. Fine adjustments to intensity thresholds are made to exclusively select APXs at the convergence points of the spindles. Employing an oversampling condition in which the laser spot size is larger than the pixel size, the treatment of centrosomes by lasers can be performed. In **Figures 6d, S11**, two centrosomes are designated for laser treatment, one with the 405 nm laser and the other with the 532 nm laser. The total laser dosage for the treatment is 0.38 mJ for the 405 nm laser and 0.41 mJ for the 532 nm laser. One hour after the treatment, a structural disruption is observed in microtubule spindles and chromosomes associated with the centrosome treated by the 405 nm laser. Following cell culture for 15 hours post-treatment (**Figures 6d, S11**), the formation of multi-nuclei was detected in the treated cells. This experiment is replicated as depicted in **Figure S11**. The fluorescence signals and bright-field transmission illumination are used to monitor the cells at different time points (**Figures S11,12**). Future studies will include extended time-lapse studies to understand the prolonged viability and productivity of the treated cells.

#### Note 8. Time-lapse imaging of EB3-EGFP cells treated with 405 nm lasers solely in the nuclei

**Figure S13** displays the time-lapse EB3-EGFP signals of HeLa cells when their nuclei are exclusively illuminated by a 405 nm laser. A longer treatment time results in a higher laser dosage within the nucleus, triggering apoptosis characterized by cell shrinkage and condensation (**Figures S13a,b**). Conversely, the treatment with a lower laser dosage does not prompt a comparable apoptotic response (**Figured S13c,d**). This investigation not only exemplifies the capability of S-RPOC in monitoring prolonged cellular reactions following precise organelle perturbation but also assesses the 405 nm laser dosage associated with nuclear treatment capable of inducing apoptosis in HeLa cells.

#### Note 9. Treatment of a single cancer cell in a co-culture system

RPOC allows for the selective treatment of cancer cells in co-culture. In this work, pancreatic cancer cells (Panc 10.05) expressing tetramethylrhodamine (TRITC) and cancer-associated fibroblasts (CAF 19) expressing fluorescein isothiocyanate (FITC) are co-cultured for selective treatment. Based on different fluorescence signals of cancer cells and fibroblasts, we outlined one cancer cell and selectively directed a 405 nm laser to interact with it without affecting a neighboring fibroblast (**Figure S14a**). The laser power for treatment is 600 μW. A substantial decrease in fluorescence signals is observed during treatment in the targeted cancer cell (**Figure S14b**). The adjacent cancer and fibroblast cells exhibit a less significant reduction in signals. Long-term imaging of cell responses reveals a recovery of fluorescent signals in the fibroblast but an irreversible loss of signals in the treated cancer cell (**Figures S14c**). Moreover, exposure to blue light prompts detachment of the treated cancer cell from its adjacent cancer cells, with less disruption for the CAF 19 (**Figures S14a,d**). These findings demonstrate distinct responses of various neighboring cells within a co-culture system to the blue-light-treated cancer cells. Further investigations into cell interactions in co-culture systems after RPOC treatment will be conducted in the future.

Supplementary Figures

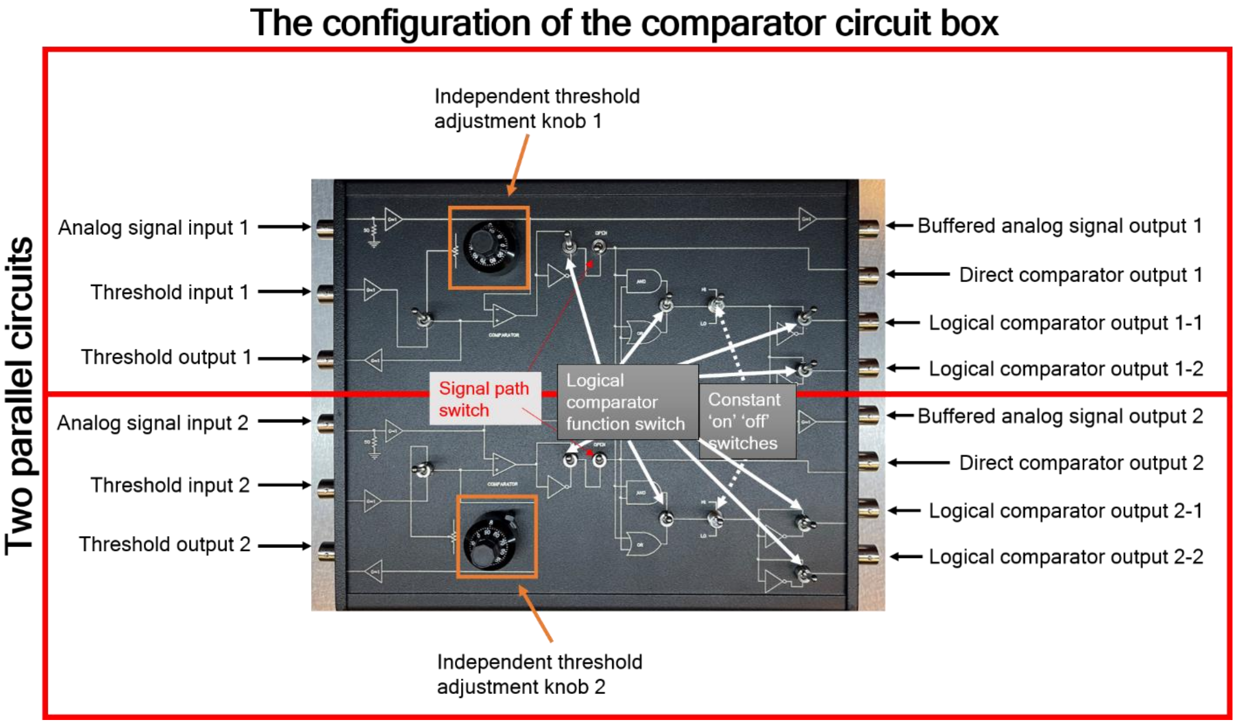


**Figure S1.** The input/output and functionalities of the dual-channel comparator circuit box.

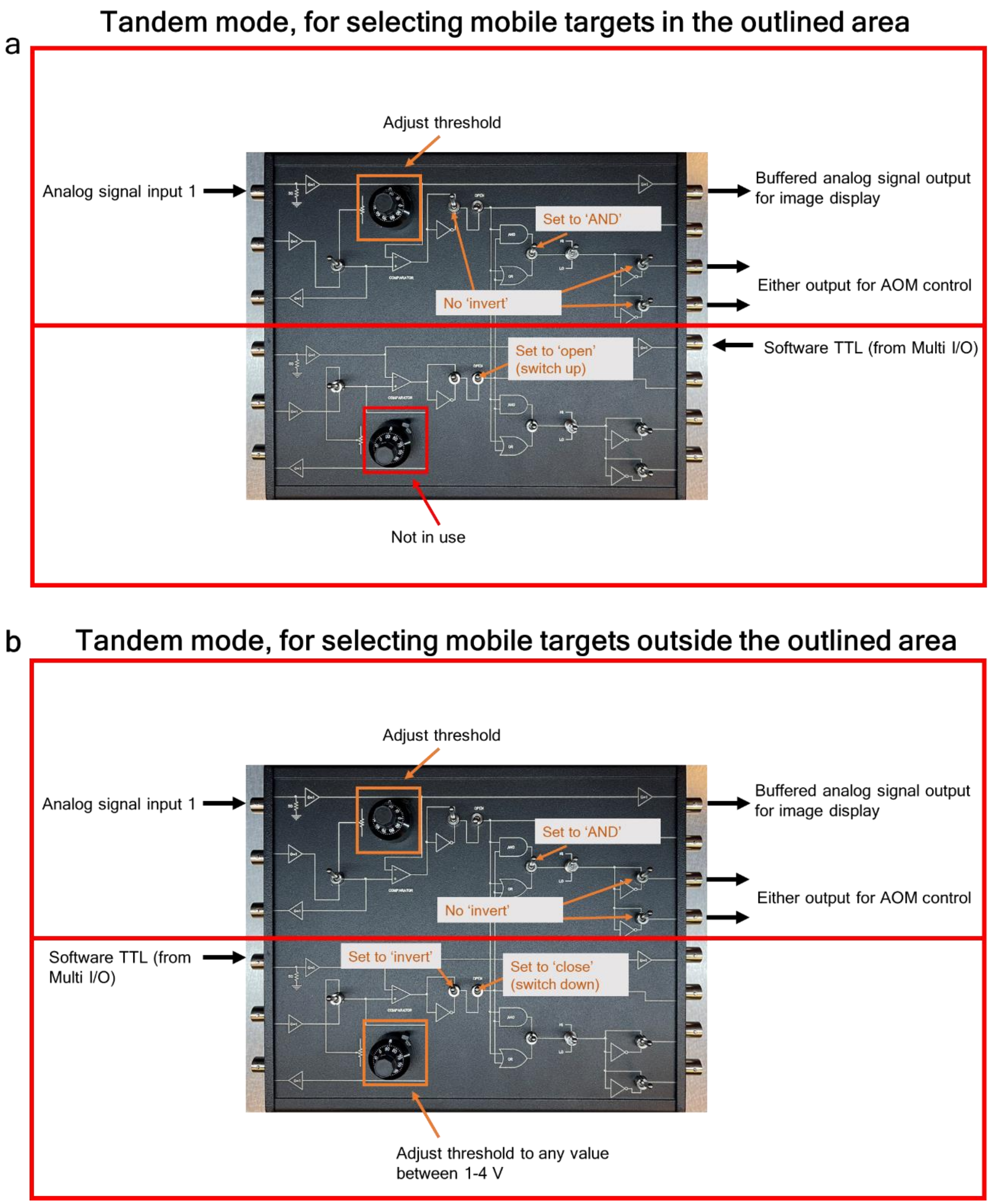


**Figure S2.** (a) One configuration of the comparator circuit box when employed in tandem with the S-RPOC software to perform real-time APX determination for mobile targets within the selected regions of interest. (b) Another configuration of the tandem mode that allows to select APXs on mobile targets outside of the outlined regions of interest. When the bottom 'invert' switch is switched to non-invert. This configuration gives the same function as panel (a).

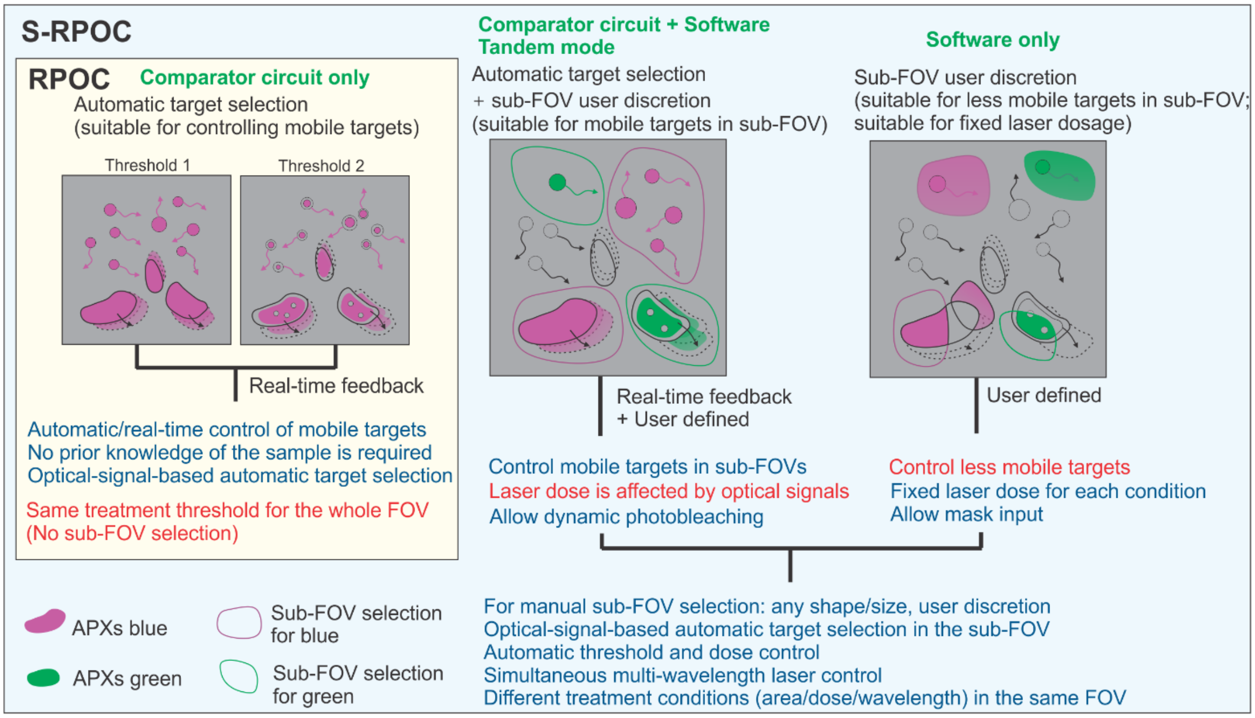


**Figure S3.** The summary of the distinctive functions and features of S-RPOC in the software-only, comparator-circuit-only, and the software + comparator-circuit tandem modes. The conventional RPOC only utilizes the comparator circuit.

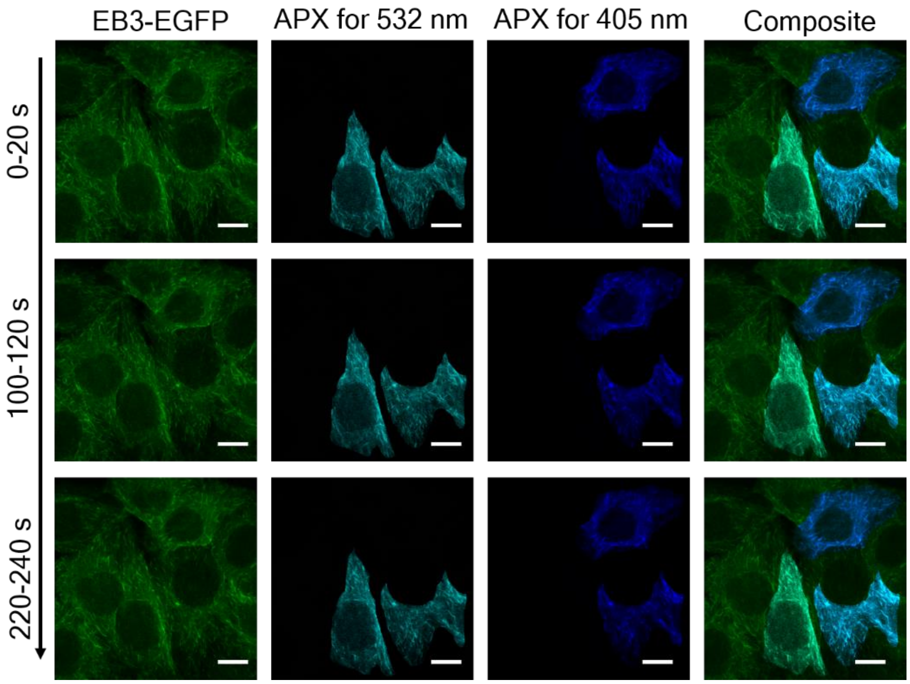


**Figures S4.** Signals from EB3-EGFP in HeLa cells and the chosen APXs based on the EB3-EGFP signals for concurrent 532 nm and 405 nm treatment. The composite images superimpose EB3-EGFP signals with all APXs. Scale bars: 10 μm.

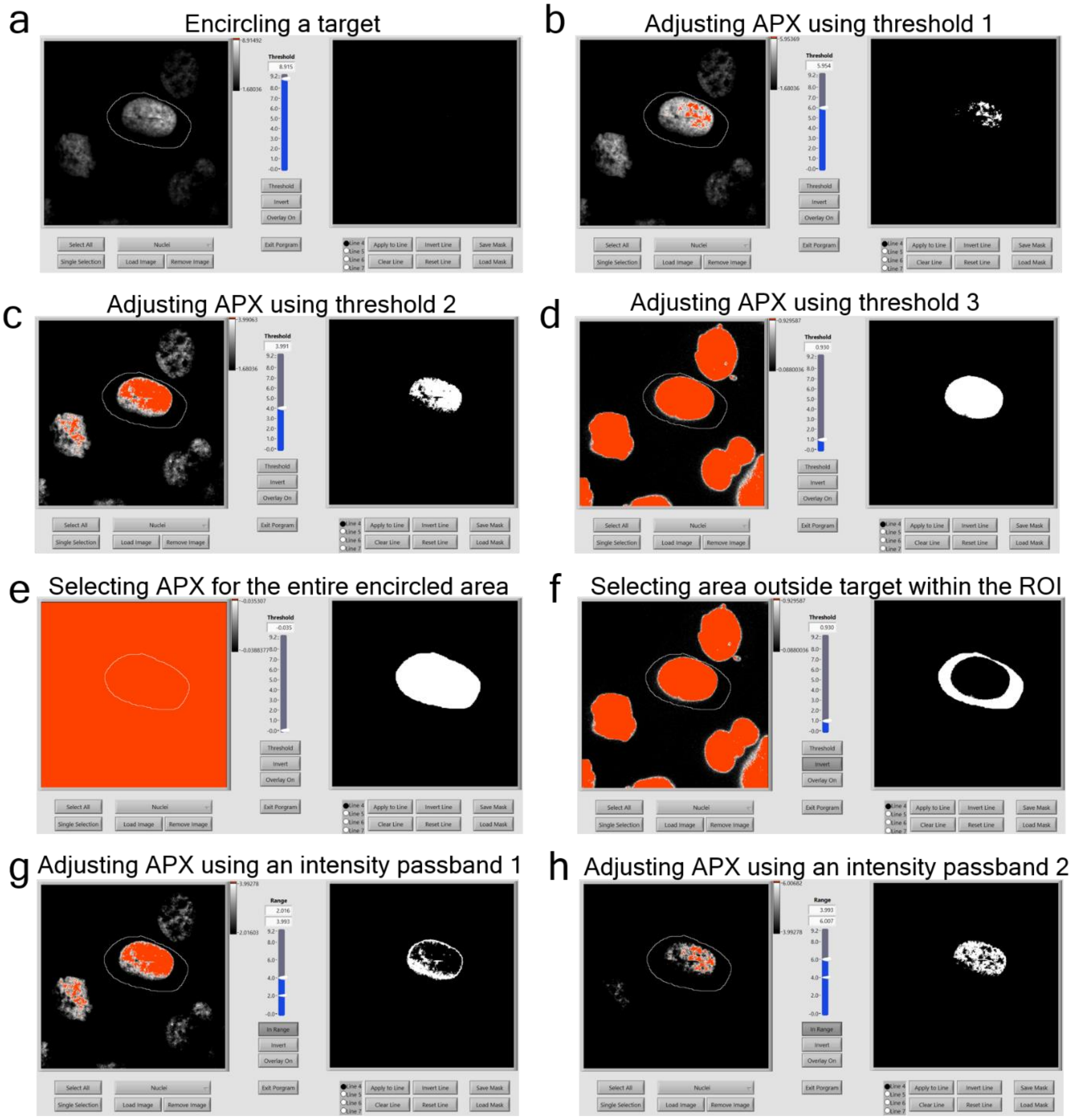


**Figure S5.** (a-d) Selection of APXs through manual outlining of a sub-FOV containing a nucleus expressing mCherry-H2B and applying different intensity thresholds. A reduced threshold value enlarges the selected nucleus area. (e) Setting the threshold to 0 V or below allows the selection of APXs that cover the entire designated area. (f) Employing the inverse function within a specific area enables the selection of APXs below the intensity threshold within the chosen region. (g,h) Utilizing the 'In Range' function facilitates the selection of APXs within a specific mCherry intensity range of the nucleus.

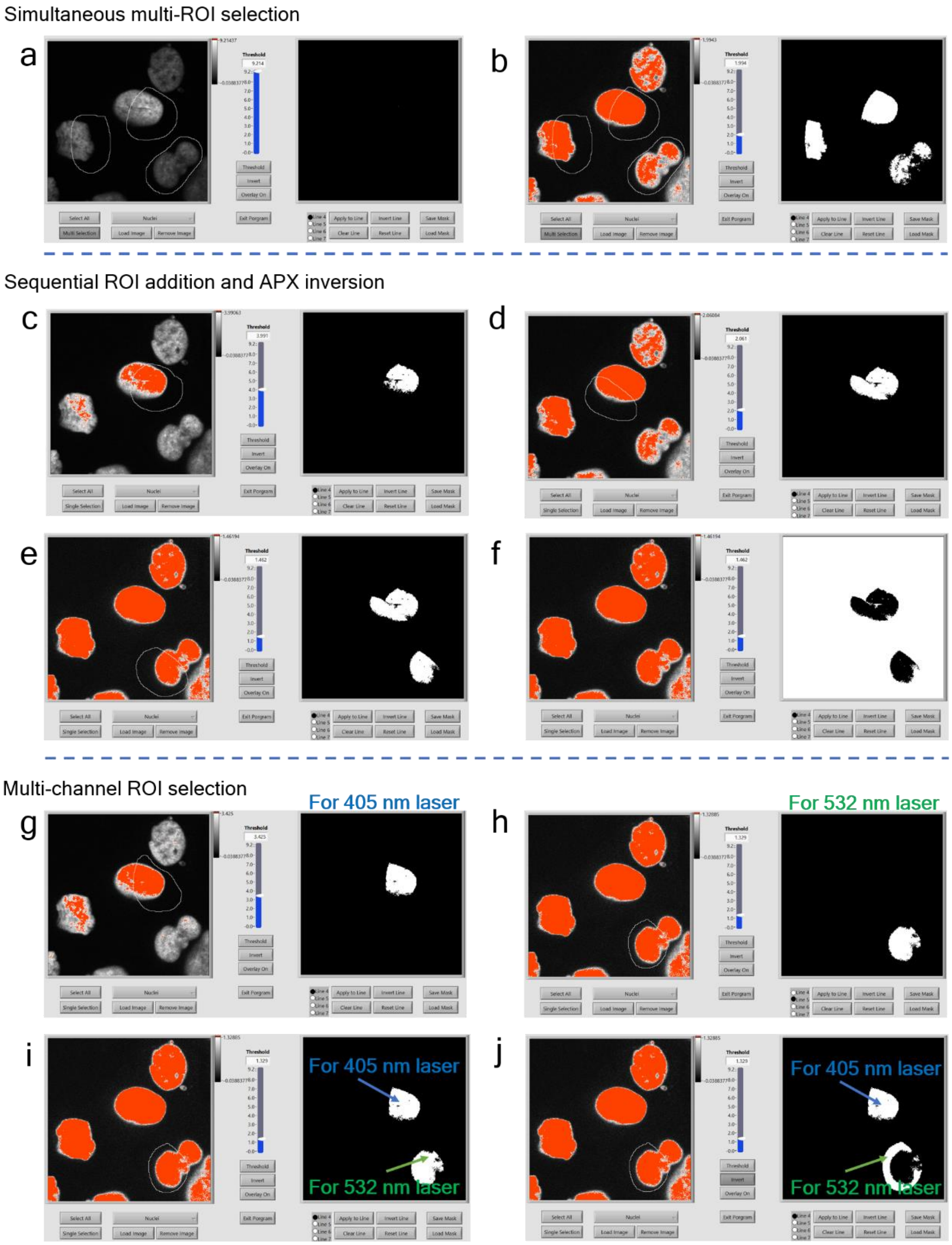


**Figure S6.** (a) Using the 'Multi-selection' function to designate multiple sub-FOVs. (b) Defining the APXs by adjusting the intensity threshold following the multi-area selection. (c-e) Sequentially adding APXs in different sub-FOVs for the 405 nm laser interaction using the 'Add to line' function. Each sub-FOV's intensity threshold can be individually adjusted. (f) Employing the 'Invert' function to reverse the selected APXs for the entire FOV after selection. (g) Designating a sub-FOV for 405 nm laser treatment. (h) Designating a sub-FOV for the 532 nm laser treatment. (i) Displaying chosen APXs for both lasers in panels (g) and (h). (j) Using the 'Invert' function exclusively for the 532 nm laser to reverse its APXs within the selected area.

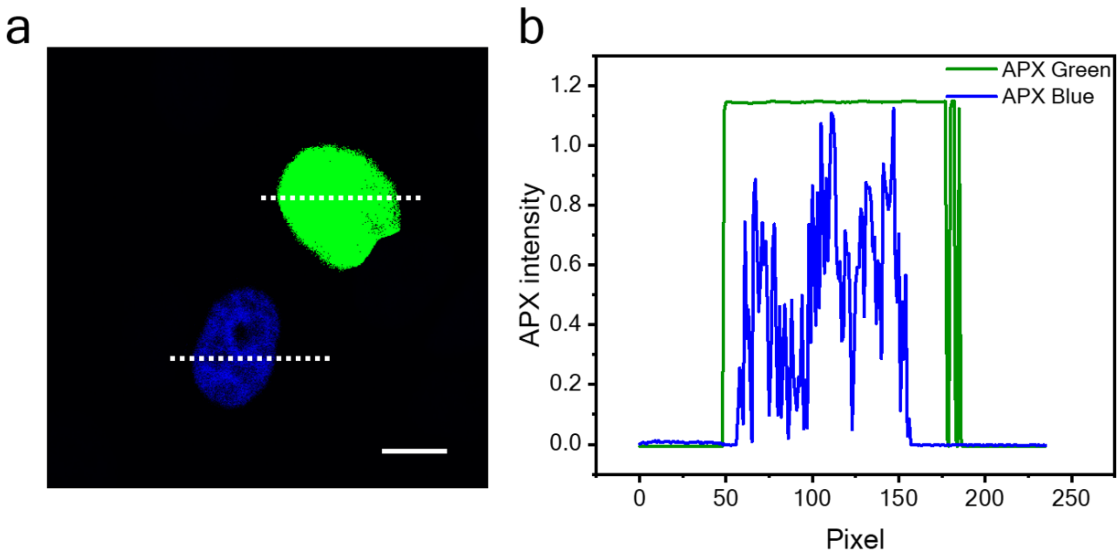


**Figure S7.** (a) The image of APXs defined only by the S-RPOC software (green) and in tandem with the software and comparator circuit (blue). (b) The corresponding intensity profiles of the white dash lines in the APX image. The scale bar is 10 μm.

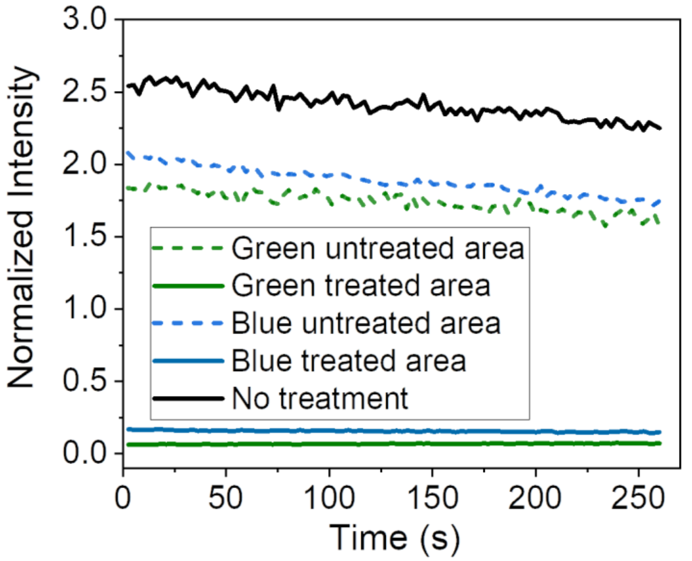


**Figure S8.** Absolute mCherry-H2B signal changes in the treated and untreated areas in the nuclei shown in Figure 4a-c.

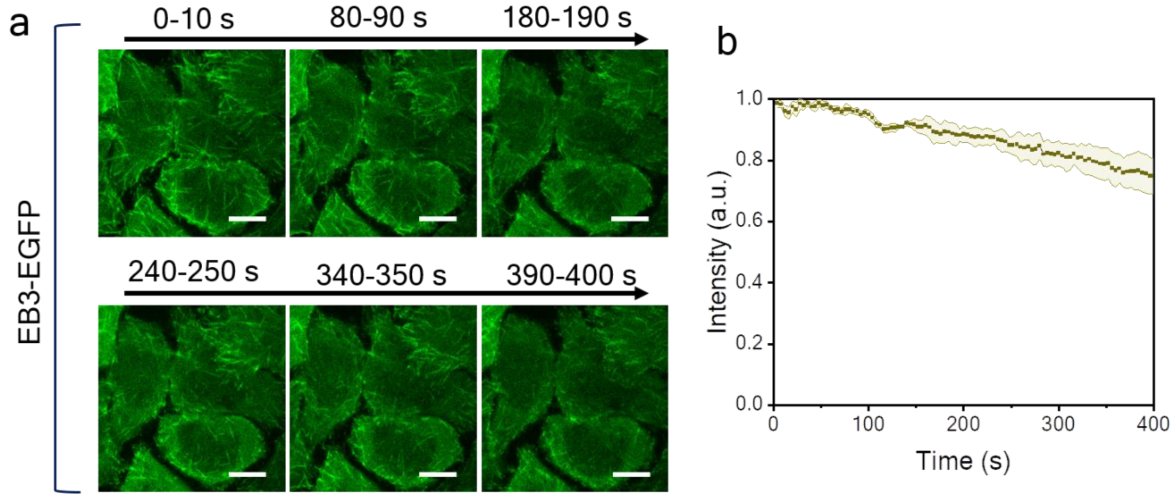


**Figure S9.** (a) EB3-EGFP signals for HeLa cells continuously excited by 25 µW 473 nm laser in different time windows. (b) Time lapse EB3-EGFP signal changes under continuous 473 nm laser scanning for 400 seconds. Scale bars: 10 µm.

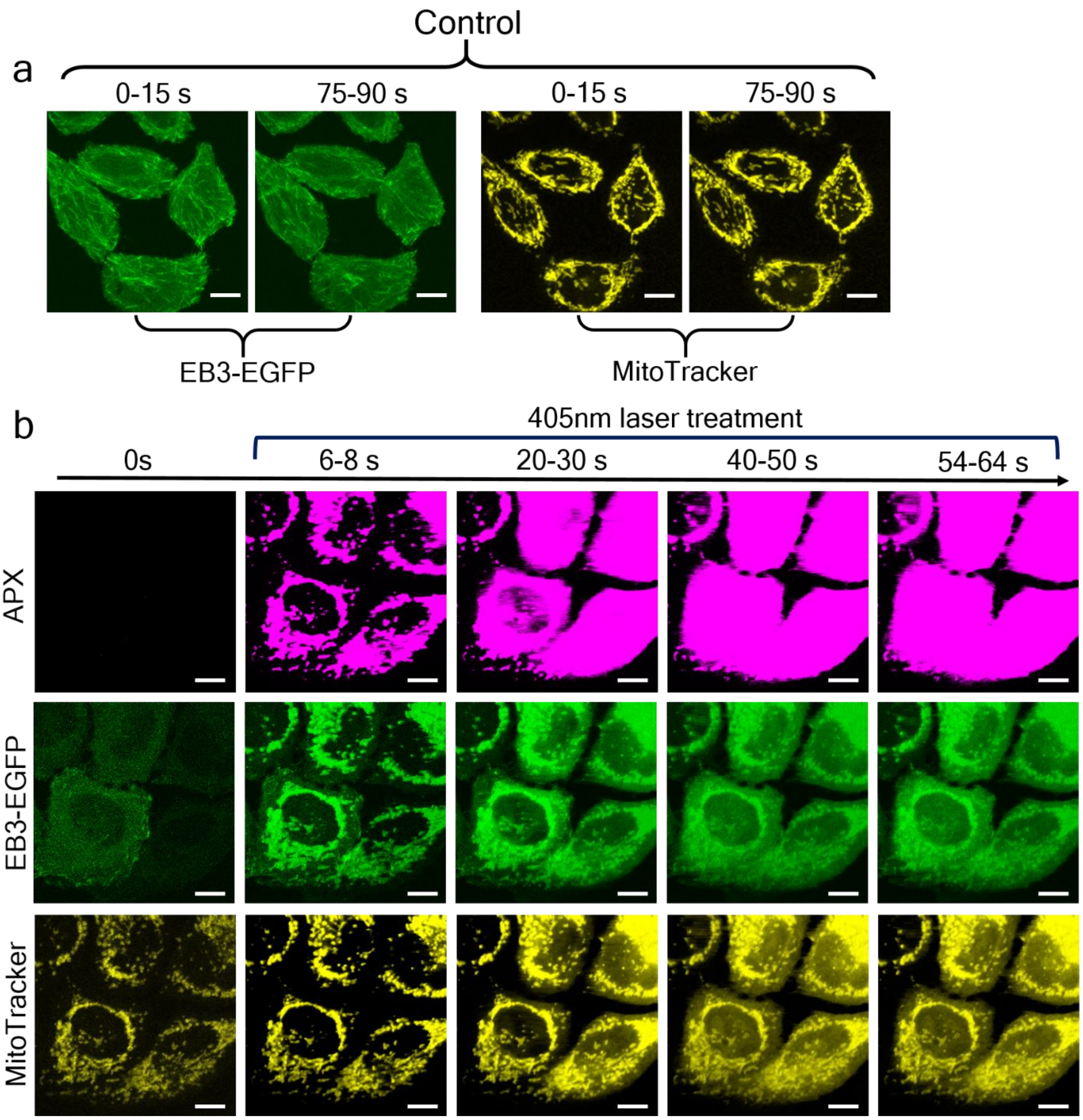


**Figure S10.** (a) EB3-EGFP (green) and MitoTracker (yellow) signals from HeLa cells continuously excited by 25 μW 473 nm laser and 10 μW 589 nm laser in two-time windows. (b) The APX (magenta), EB3-EGFP (green), and MitoTracker (yellow) signals during 405 nm laser treatment targeting mitochondria. Here, only the comparator circuit is used to determine APXs. The S-RPOC software is not applied. Scale bars: 10 μm.

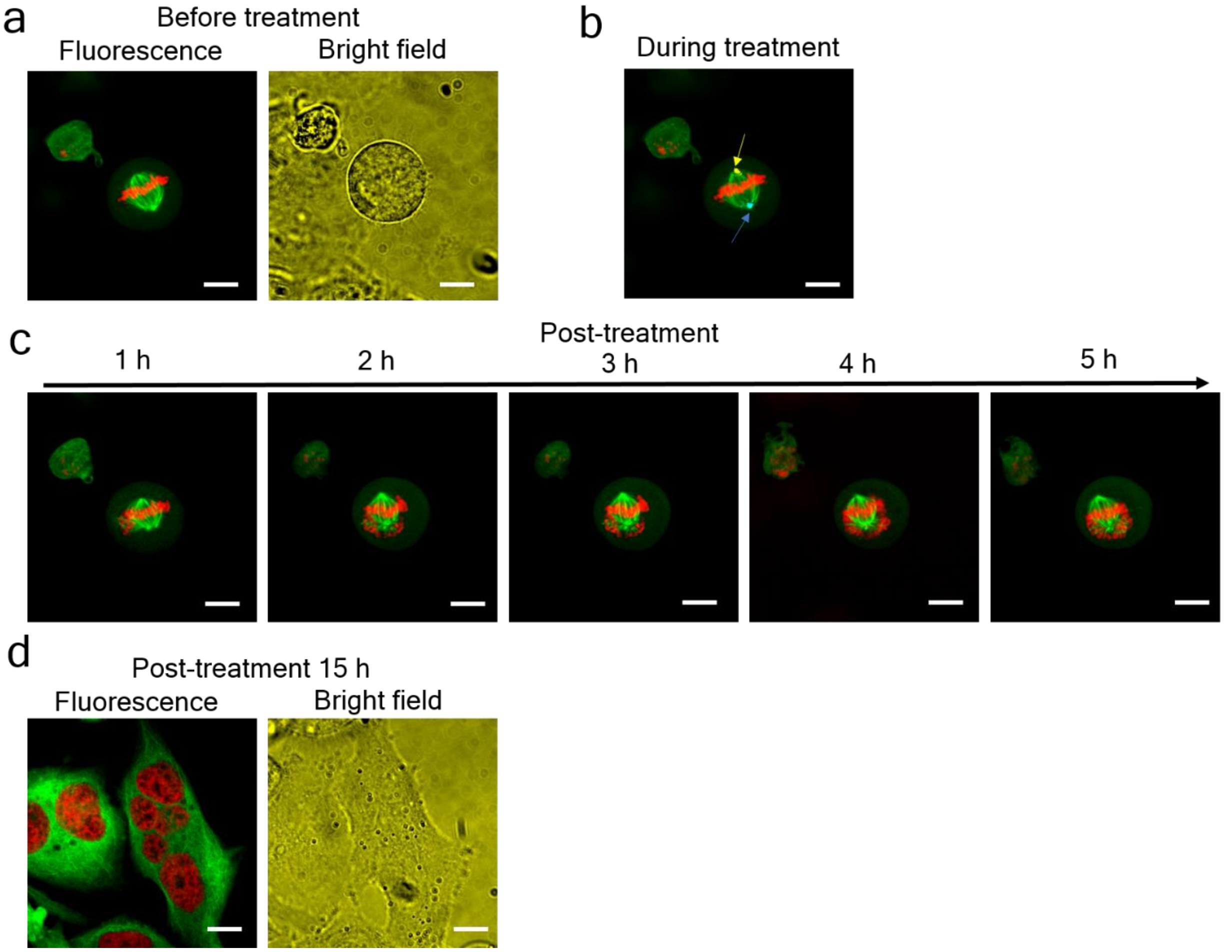


**Figure S11** (a) A HeLa cell expressing mCherry-H2B (red) and EGFP-tubulin (green) in the mitotic phase visualized using the S-RPOC fluorescence imaging channels and the bright-field transmission mode. (b) Treatment of centrosomes using 240 μW 405 nm (blue) and 240 μW 532 nm (yellow) lasers for 275 seconds. (c) Time-lapse images of the treated cell at different time points after treatment. (d) After 15 hours, the cell transitions into a multinucleated form. Scale bars: 10 μm.

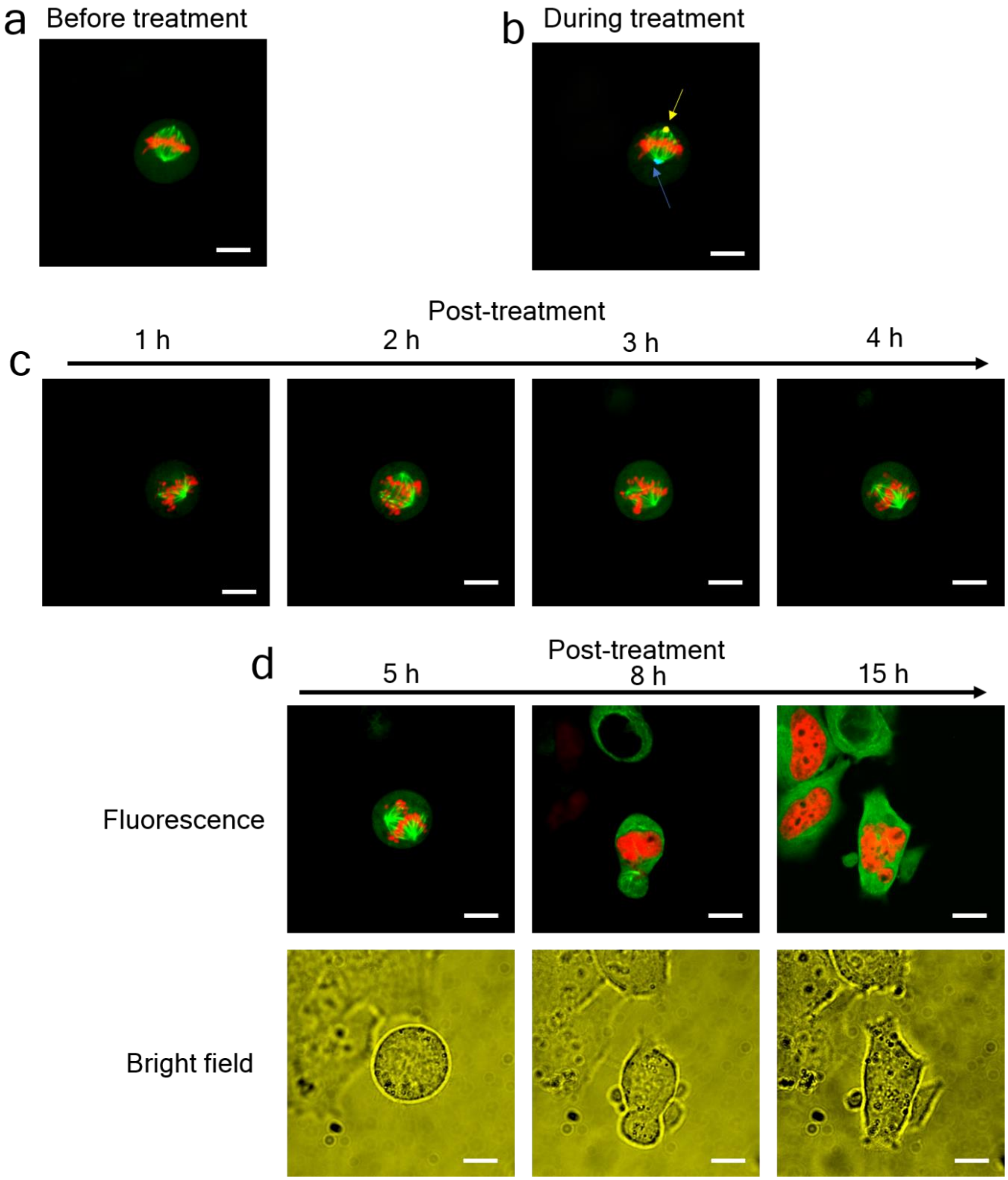


**Figure S12 Replicating the modulation of cell division using S-RPOC via the treatment of centrosomes.** (a) A HeLa cell expressing mCherry-H2B (red) and EGFP-tubulin (green) in the mitotic phase visualized using the S-RPOC fluorescence imaging channel. (b) Treatment of centrosomes using 240 μW 405 nm (blue) and 240 μW 532 nm (yellow) lasers. (c) Time-lapse images of the treated cell at different time points post-treatment. (d) Time-lapse fluorescence and bright-field images of the treated cell at different time points after 5 hours. Scale bars: 10 μm.

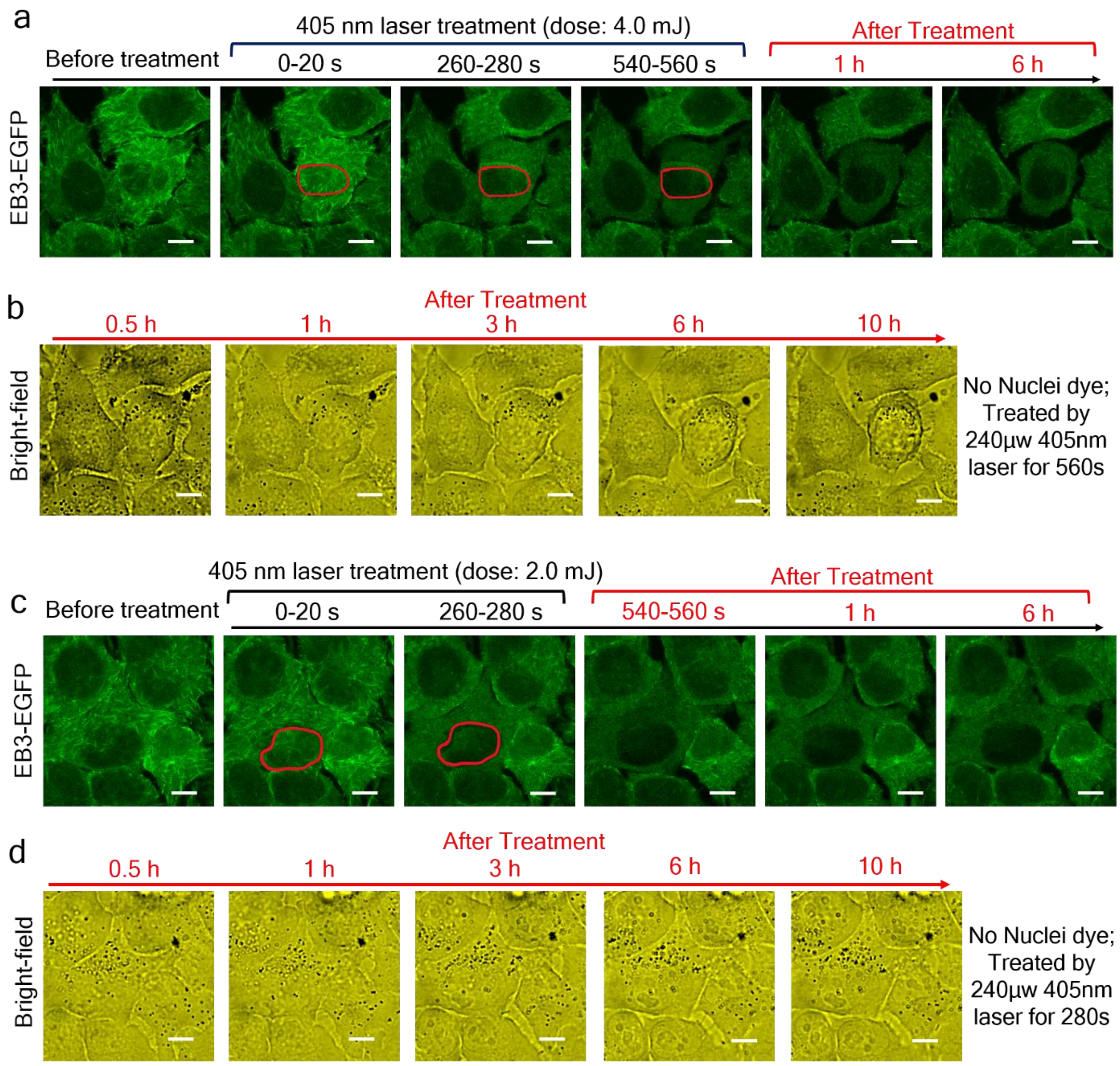


**Figure S13** (a) EB3-EGFP signals from HeLa cells before, during, and after treatment of the nucleus of a specific cell by using a 240 μW 405 nm laser. The unlabeled nucleus is chosen based on the contrast of the EB3-EGFP signals. (b) Bright-field images of the same FOV in panel (a) at different time points after treatment. (c) Similar to panel (a), exhibiting EB3-EGFP signals, but treated with only 240 μW 405 nm laser. (d) Similar to panel (b), illustrating the same FOV and cells featured in panel (c) using bright field imaging. Scale bars: 10 μm.

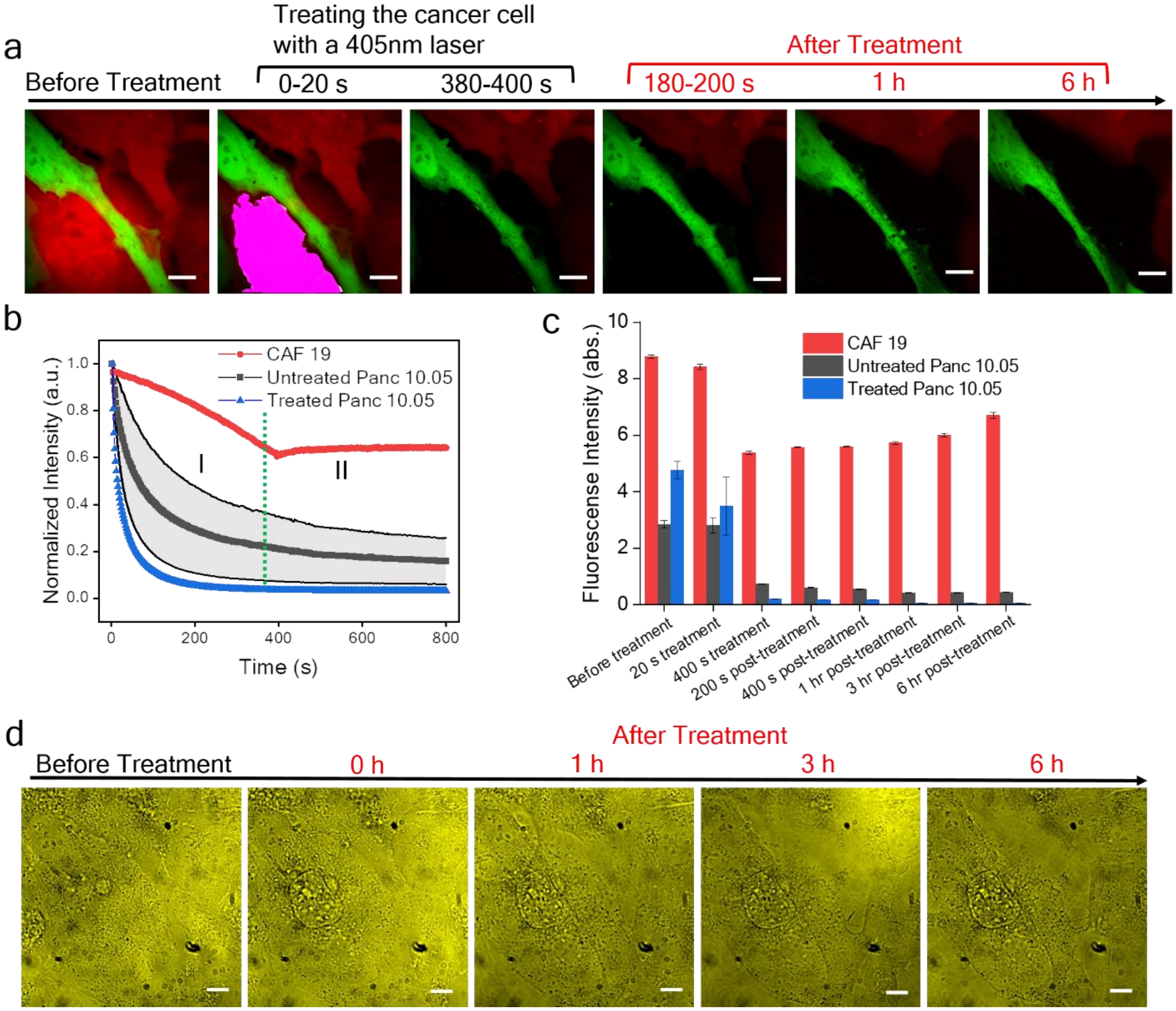


**Figure S14** (a) Time-lapse fluorescence images of Panc. 10.05 cells (red) and CAF 19 cells (green) before, during, and after treatment with a 240 μW 405nm laser targeting the selected APX highlighted in magenta. (b) Temporal changes in fluorescence signals for the CAF 19 cell (red), the untreated Panc. 10.05 cell (black), and the treated Panc. 10.05 cell (blue) during treatment and the subsequent 400 seconds. (c) Post-treatment, the fluorescence signals from CAF 19 cells (red columns) notably decrease but gradually recover in a 6-hour duration. Both untreated (black columns) and treated Panc. 10.05 cells exhibit decreased fluorescence signals, with a more pronounced reduction observed in the treated cell. (c) Bright-field images captured before and after treatment within the same FOV at different time points. The detachment of the treated cell from neighboring cells and the condensation of the nucleus in the treated cell are detected. Scale bars: 10 μm.

### Supplementary Videos

Video S1. Adaptive designation of laser interaction regions by S-RPOC

Video S2. Controlling laser interactions through mask input

Video S3. Selective laser interactions using S-RPOC targeting mCherry-H2B within nuclei of live cells.

Video S4. EB3-EGFP signals in HeLa cells and the chosen APXs based on the EB3-EGFP signals for concurrent 532 nm and 405 nm laser treatment.
